# Discovery of frameshift-associated TCRs following personalized neoantigen vaccination in a patient with osteosarcoma

**DOI:** 10.64898/2026.10.09.757920

**Authors:** Sarah N. Bennett, Jessica Alley, Mitra Shabrang, Jason Garness, Karen McKinnon, Amber Smith, William B. Inabinett, William A. Hoos, Marshall A. Thompson, Alex Rubinsteyn, Benjamin G. Vincent

**Author notes:** Corresponding Author:* Benjamin G. Vincent, MD, Associate Professor of Medicine, Division of Hematology, Joint Faculty, Department of Microbiology and Immunology, Curriculum in Bioinformatics and Computational Biology, Computational Medicine Program, Lineberger Comprehensive Cancer Center, University of North Carolina at Chapel Hill, 5206 Marsico Hall, Chapel Hill, NC 27599, @BenjaminGVincen.

## Abstract

Personalized antigen-directed immunotherapies, including personalized cancer vaccines, are emerging as promising therapeutic modalities for a range of cancer types. Immune monitoring can be used to evaluate the efficacy of a vaccine in stimulating the immune response pre- and post-vaccination; however, such approaches typically stop short of identifying which T cell receptors (TCRs) are functionally driving therapeutic responses. Here, we present a case report of immune monitoring and subsequent TCR discovery experiments following administration of personalized cancer vaccines in a patient with locally recurrent osteosarcoma. Whole PBMCs were stimulated with vaccine peptides (>16 AA) *in vitro* for 11 days, after which IFNγ ELISPOT identified total T cell responses for 12 of 15 tested peptides post-vaccination. *In vitro* expanded samples were submitted for 10x single-cell TCR sequencing, from which 26 TCRs were chosen for generation of TCR-T cells based on clonal expansion and culture specificity. Of the generated TCR-T cells, 25 produced functional TCRs, 4 of which were specific to vaccine peptides. However, downstream testing revealed each was MHC class-II restricted. To address this, we repeated the *in vitro* stimulations with co-cultures of isolated CD8^+^ T cells with autologous dendritic cells, and cultures of whole PBMCs with anti-MHC class-II blocking antibody. These cultures yielded 10 additional TCRs, 2 of which were responsive to a vaccine peptide containing a MAP2 frameshift mutation. These findings demonstrate an effective approach for leveraging post-vaccination expanded antigen-specific T cell populations for discovery of novel T cell receptors for downstream cellular therapies.

**Graphical Abstract:** 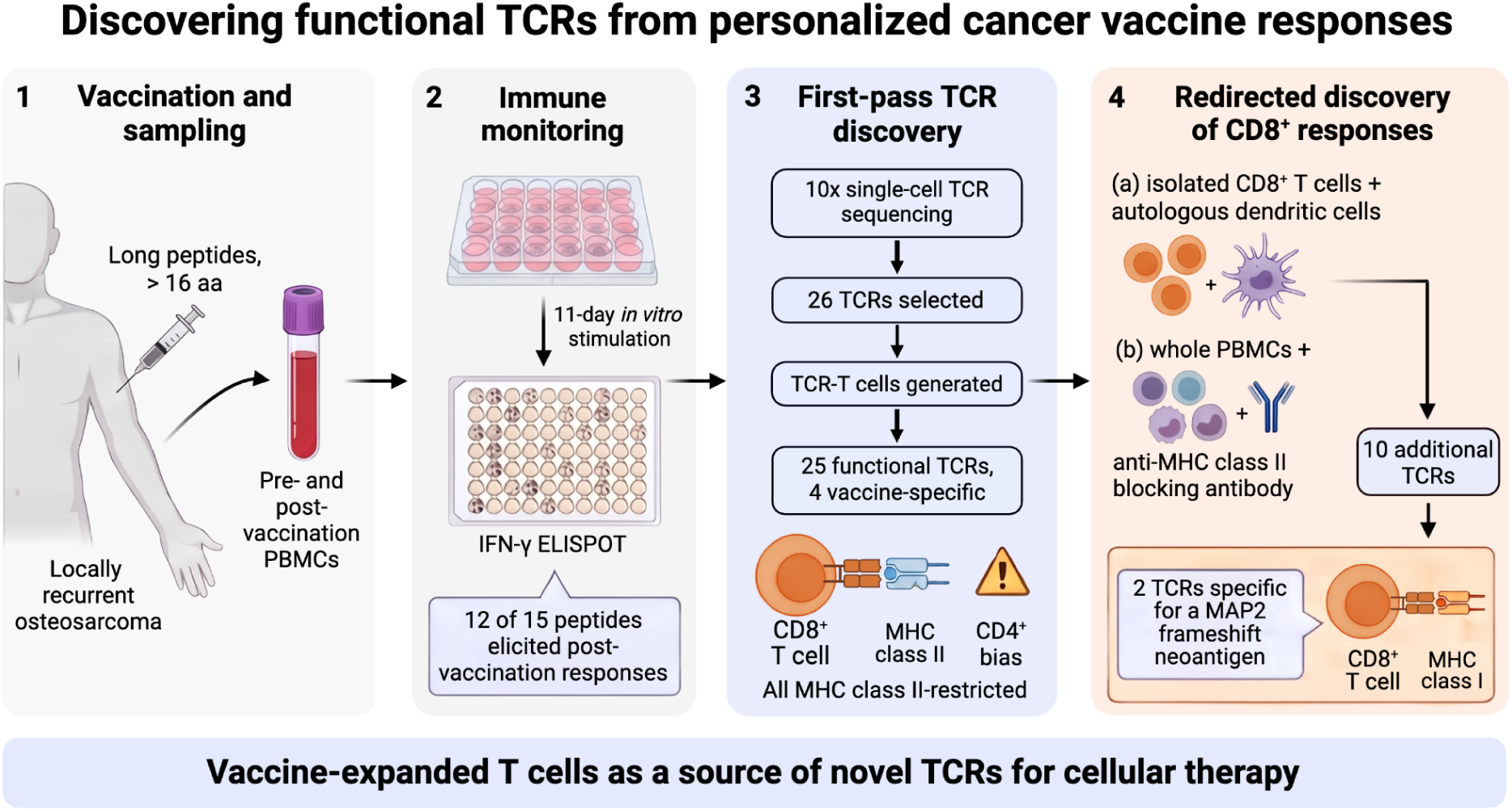

## Introduction

Personalized antigen-directed immunotherapies targeting peptide antigens presented by HLA molecules are emerging as promising cancer therapies. Modalities include personalized cancer vaccines (PCVs)^1,2^, endogenous T cell therapy (ETC)^3–5^, antigen-specific T cell selected tumor-infiltrating lymphocyte therapy (SEL-TIL)^6,7^, and T cell receptor engineered T cell therapy (TCR-T)^8–10^. Therapeutic cancer vaccines, including peptide and mRNA formulations, have had success in clinical trials^11–16^, and ETC, SEL-TIL, and TCR-T approaches have yielded clinical responses in small cohort studies^5–7,17–21^. Each approach requires selection of potentially immunogenic tumor antigens, most commonly done by analyzing DNA and/or RNA sequencing data for tumor-specific variants. Additional selection criteria commonly include *in silico* binding predictions of a given peptide to the patient’s specific HLA proteoforms^22,23^. Although still largely underutilized due to significant barriers in cost and experimental design, the highest standards for antigen verification include experimental validation of peptide presentation by mass spectrometry, and/or immunogenicity using patient-derived T cell assays^24,25^.

Tumor antigens can be encoded by multiple classes of genomic sources, including (1) somatic mutations such as single nucleotide variants (SNVs), insertion/deletion variants, frameshift variants, and gene fusions, (2) expressional antigens that are encoded in the germline DNA but expressed selectively in tumor cells, such as cancer/testis antigens, endogenous retroviral antigens, and tumor-specific alternative splice variants, (3) tumor-specific protein degradation products, and (4) viral genes expressed by oncogenic viruses^26^. Tumor antigens arising from somatic mutations have become known as “neoantigens,” with SNVs representing the most frequently investigated source. These neoantigens have the advantage of being relatively easy to identify from clinical short-read sequencing assays, and a fraction of these, including key examples derived from oncogenic mutations at KRAS G12 and PIK3CA H1047, have been found to be immunogenic *in vivo*^18,27^. Despite these successes, the majority of SNVs do not elicit immunogenic neoantigens, and most predicted neoantigens derived from SNVs are not presented by HLA and/or are not immunogenic^28^. There is thus a need to consider neoantigen sources beyond SNVs to expand the set of targetable tumor antigens. Frameshift mutations are especially attractive for this purpose, as they are demonstrably tumor-specific compared to expressional antigens, but also elicit peptides with longer stretches of “non-self” amino acids than SNV-derived neoantigens. The importance of targeting frameshift mutations is becoming increasingly apparent, and several groups have recently published characterizations of T cell responses, supporting their potential for use in targeted cellular therapies^29,30^.

TCR-T therapy requires identification and characterization of antigen-specific TCRs for use as a cellular therapy product. TCRs derived from CD8^+^ T cells are commonly prioritized for their cytotoxic potential and ability to recognize antigens presented by MHC-I, as opposed to TCRs derived from CD4^+^ T cells, which are associated with helper functions and MHC-II antigen recognition. Following discovery of TCRs specific for target tumor antigens, candidate TCRs may be prioritized for inclusion in therapeutic products based on features including functional avidity, predicted peptide/HLA target density, and cytotoxicity against tumor cells or other antigen-presenting target cells^10,25,31^. TCR-T based-therapies have several theoretical advantages over other personalized antigen-directed therapies, including knowledge of and control over the specific TCR(s) used, capacity to benchmark and select TCRs for inclusion based on functional activity and target distribution across HLA proteoforms, and ability to achieve higher cell doses per TCR for a small set of highly active TCRs, compared to more polyclonal populations in ETC and TIL therapies.

However, TCR-T therapies are limited by the difficulty of discovering antigen-specific TCRs in any given patient. Antigen-specific T cell populations often exist at low frequencies in circulation, with discovery often requiring capture methods capable of isolating less than a single cell in a million^32^. TCRs have been successfully isolated from tumor-infiltrating lymphocytes (TILs)^18,33,34^ and from naïve cell priming of healthy donor cells^35^, although debate persists as to the advantages of one approach compared to the next. For either approach, the historical gold standard has been affinity-based isolation using peptide-MHC multimers. However, lack of commercial reagent availability for all but the most common of HLA alleles, coupled with challenges arising from in-house generation of peptide-MHC monomers, restricts TCR discovery efforts. As such, a range of function-based isolation approaches has emerged, including selection based on cytokine production^36^, expression of activation markers^37,38^, or clonal expansion, assessed either *in vitro* or bioinformatically^39^. The success rate of these approaches at identifying TCRs of interest may be further improved by strategies to increase the overall frequency of antigen-specific T cells *in vivo*, prior to any method of *in vitro* expansion.

Here, we report a series of tumor antigen-specific TCR discovery experiments following administration of three rounds of personalized neoantigen vaccines in a patient with locally recurrent osteosarcoma. We corroborate previous reports that vaccination with long peptides (>16 AA) and subsequent standard immune monitoring approaches are biased towards priming and detection of *ex vivo* CD4^+^ T cell responses^2,40–42^, and provide an alternate approach to detect CD8^+^ T cell responses with the ultimate goal of discovering and validating therapeutically relevant TCRs. Using this approach, we discovered a pair of novel frameshift mutation-specific TCRs recognizing a minimal epitope included in the original set of neoantigen vaccine peptides.

## Materials and Methods

### Study Design

The aim of this study was to evaluate if endogenous immune responses against tumor antigens are boosted post-neoantigen vaccination, and ultimately, to isolate clinically relevant TCRs for potential use in a TCR-T therapy. To this end, pre- and post-vaccine PBMC samples were collected under RCRF research protocols H37001 and H37002. Whole exome sequencing and details pertaining to vaccine design, manufacturing, and administration were performed externally by the Jamie Leandro Foundation in two formulations, referred to here as JLFv1 and JLFv2. In both formulations, synthetic long peptides containing both MHC-I and MHC-II epitopes were delivered as injections accompanied by a poly-ICLC adjuvant, given as a series of priming doses followed by up to four boosts. For use in immune monitoring and downstream assays, peptides were sourced directly from the JLF vaccine manufacturer, CSBio (Menlo Park, CA). Additional deidentified clinical metadata associated with vaccination and clinical samples were also provided.

### PBMC Stimulation and Culture

Pre- and post-vaccine PBMC samples were viably thawed and plated at 5 × 10^6 cells/mL in a 24-well untreated cell culture plate in RPMI supplemented with nonessential amino acids, HEPES, L-glutamine, sodium pyruvate, penicillin/streptomycin, B-mercaptoethanol, and 10% heat-inactivated, AB-positive human serum (Sigma-Aldrich)^41^. Individual cultures were stimulated with pooled (pre-vaccine) or individual (post-vaccine) vaccine peptides at 10 ug/mL in the presence of IL-7 (20 ng/mL) (StemCell). Peptides were sourced from the vaccine manufacturer, CSBio (Menlo Park, CA) and resuspended in DMSO. Every three days, one half of the media volume was refreshed, and cultures were supplemented with additional IL-7 (20 ng/mL) and IL-2 (20 U/mL) (StemCell). Both pre- and post-vaccine timepoints included an unstimulated, negative control well; the post-vaccine timepoint also included a positive control well stimulated with PepTivator CEF viral peptide pool (Miltenyi Biotech). Both samples were stimulated with peptides from the JLFv1 vaccine formulation (n=9 peptides), while only the post-vaccine sample received stimulation with additional JLFv2 peptides (n=6 additional peptides). To comprehensively profile the post-expansion TCR repertoire, cells from all fifteen post-vaccine peptide-stimulated cells were harvested on day 11 of culture and sent for single cell TCR alpha/beta sequencing using the 10x Genomics 5’ TCR V(D)J protocol.

### PBMC ELISPOTs

On day 11 of culture, up to three replicates of 2 × 10^5 cells were restimulated with respective individual peptides at 10 ug/mL and transferred to a pre-coated IFNγ ELISPOT plate (BD Biosciences) for overnight incubation. Plates were developed according to the manufacturer’s protocols using the AEC substrate kit (BD Biosciences). Due to limiting cell numbers in the pre-vaccine timepoint, only a single replicate of restimulated cells was included. For post-vaccine samples, both restimulated and DMSO controls were included in triplicate for each individual peptide stimulated culture. All samples included PHA and PepTivator CEF viral peptide positive controls (Miltenyi Biotech). Developed plates were allowed to dry completely before being read for spot count and activity on an AID ELISPOT Plate Reader. Spot counts were normalized to the DMSO-treated co-culture negative control.

### Co-culture of CD8^+^ T cells + Autologous DCs

To generate autologous dendritic cells (DCs), monocytes were isolated from total PBMCs using plastic adherence. Briefly, PBMCs were plated at 7.5e6 cells/mL and allowed to adhere to tissue culture-treated flasks for 90 minutes, followed by vigorous washing and a second 60 minute incubation. For differentiation into immature DCs, adherent cells were cultured in RPMI containing 5% human serum, supplemented with 1,000 IU/mL GM-CSF (StemCell) and 1,000 IU/mL IL-4 (StemCell). After 5 days, immature DCs were matured overnight with 10 ng/mL LPS (eBiosciences) and 100 IU/mL IFNγ (StemCell). Finally, DCs were peptide-pulsed and cultured with 10 ug/mL peptide, 2,000 IU/mL IL-4, 572 IU/mL GM-CSF, and 20 ng/mL TNFα (StemCell). Mature DCs were harvested on ice and co-cultured at a 4:1 T cell to DC ratio with CD8^+^ T cells isolated via negative selection (StemCell) in the presence of 20 ng/mL IL-7. Media and cytokines were refreshed every three days, with the addition of IL-2 (20 U/mL, StemCell) beginning on day 3. On day 11, cultures were harvested and sent for single cell TCR alpha/beta sequencing using the 10x Genomics 5’ TCR V(D)J protocol.

### 10x Single Cell TCR Sequencing and Analysis

For selected samples, 10x Genomics Single Cell Sequencing was performed using the 5’ TCR V(D)J protocol via the UNC Immune Monitoring and Genomics Facility (IMGF) and the UNC High-Throughput Sequencing Facility (HTSF). Selection of TCR sequences for further analysis was initiated with the 10x Genomics CellRanger VDJ pipeline. Cells were then filtered to contain at least 200 detected genes, at least 1,000 UMIs, and under 20 percent mitochondrial reads. GEX data was incorporated to assign cells as CD8^+^ T cells, CD4 T^+^ cells, or unknown, based on expression of CD8 alpha, CD8 beta, and CD4 gene signatures. Sequences were then grouped by CDR3 clonotype and filtered by frequency and overlap between stimulation cultures. Remaining clonotypes were matched against publicly available TCR databases, including VDJdb^43^, IEDB^44^, and CEDAR^45^, and those with known specificities were eliminated. Finally, twenty-six full length TCR sequences with beta-T2A linker-alpha orientation were generated.

### Generation of TCR-T Cells

TCR-T cells were generated as previously described by knocking-out endogenous alpha and beta TCR chains, followed by a gamma-retroviral transduction to express novel TCRs of interest^9,10^. Briefly, healthy donor PBMCs were sourced from Gulf Coast Blood, isolated via Ficoll-Paque (Cytiva) density gradient separation, and viably cryopreserved in Cryostor-CS10 (StemCell) until use. After viably thawing, total CD8^+^ T cells were negatively selected with an EasySep Human CD8^+^ T Cell Isolation Kit (StemCell) and stimulated for 72-hours using plate-bound anti-CD3 and anti-CD28 (1 ug/mL, BioLegend). During activation, CD8^+^ T cells were cultured with IL-2 (300 IU/mL, StemCell), IL-7 (5 ng/mL, StemCell), and IL-15 (5 ng/mL, StemCell). Following activation, T cells were harvested, washed in PBS, and electroporated (pulse code: EH115) with ribonucleoproteins containing guide-RNAs targeting the TCR alpha and beta chains using a 4D Nucleofector X (Lonza)^46^. Cells were cultured for 48-72 hours with 180 IU/mL IL-2 before retroviral transduction.

Retroviral supernatants were generated by transfecting HEK293T cells with retroviral plasmids containing the novel TCR of interest. Cells were transfected using GeneJuice Transfection Reagent (Sigma-Aldrich), according to the manufacturer’s protocol, and supernatants were harvested at 48- and 72-hours post transfections. TCR knock-out cells were then transduced in RetroNectin-coated (TakaraBio) plates. Non-treated 24-well plates were coated overnight at 4℃. Prior to adding virus, plates were blocked with complete media for 10 minutes at room temperature. 2 mLs of virus was added per well, and plates were spun for 90 min at 2000 *x g*. Supernatant was aspirated and 2 × 10^5 TCR knock-out cells were added at 1 × 10^5/mL with 10 ng/mL IL-7 and IL-15 (StemCell). Cells were spun for 10 minutes at 1000 *x g*, then incubated at 72℃ for a minimum of 72 hours before screening.

### Validation of TCR Expression by Flow Cytometry

To confirm successful editing of TCR-T cells, samples were stained on ice with monoclonal antibodies targeting PE-TCRαβ (1:200; IP26, BioLegend) and BV421-CD8 (1:400; SK1, BioLegend) or BUV395-CD8 (1:400; RPA-T8, BD Biosciences). Cells were also stained with fixable NIR viability dye (1:2,000; ThermoFisher), run on a BD Symphony A3 (BD Biosciences), and analyzed in FlowJo V10.10.0.

### TCR-T ELISPOTs

TCR-T specificity was evaluated via IFNγ ELISPOT (BD Biosciences) using patient-derived Epstein-Barr virus transformed T cells (EBV-LCLs), which are capable of presenting both MHC-I and MHC-II peptides, as antigen-presenting cells. EBV-LCLs were generated using a previously established protocol^47^ and pulsed with 10 ug/mL peptides or DMSO overnight before use. All ELISPOTs were performed using a 10:1 effector-to-target ratio and up to 50,000 TCR-T cells per well. Each TCR-T was tested in duplicate against predicted peptide(s) and DMSO controls. Where relevant, minimal epitopes or wild-type long peptides were custom ordered at >95% purity from GenScript and resuspended in DMSO. Plates were incubated overnight at 37℃ and developed as described above. For determination of class-I or class-II reactivity, ELISPOTs were conducted with 30 ug/mL MHC-II blocking antibody, anti-human HLA-DR, DP, DQ (BD Biosciences).

### Intracellular Cytokine Staining

To stimulate cytokine production, TCR-T cells were co-cultured with DMSO or peptide-pulsed autologous EBV-LCLs for 6 hours in the presence of Monensin and Brefeldin A. Cells were then stained on ice with the following: BUV395-CD8 (1:400; RPA-T8, BD Biosciences), BV421-CD4 (1:400; OKT4, BioLegend), BUV737-CD45RA (1:50; HI100, BD Biosciences), Super Bright 780-CCR7 (1:50; 3D12, ThermoFisher), and PE-TCRαβ (1:200; IP26, BioLegend), as well as fixable NIR viability dye (1:2000; ThermoFisher). Samples were fixed and permeabilized using the BD Cytofix/Cytoperm Fixation/Permeabilization Kit (BD Biosciences) according to the manufacturer’s instructions before overnight staining at 4℃ with additional antibodies: BV711-IFNγ (1:200; B27, BioLegend), Alexa Fluor 647-TNFα (1:200; Mab11, BioLegend), and PE-Cy7-IL-2 (1:100; MQ1-17H12, BioLegend). Cells were fixed, run on a Cytek Aurora (Cytek Biosciences), and analyzed in FlowJo V10.10.0.

## Results

### Enhanced T cell responses following neoantigen vaccination are detected by IFNγ ELISPOT

We first aimed to understand the specificity of the immune response following personalized vaccination with predicted neoantigen long peptides via *in vitro* stimulation of total PBMCs followed by IFNγ ELISPOT (Figure 1A). For this purpose, we compared pre-vaccine and post-vaccine samples, collected approximately 8 months apart. During this period, the patient received three courses of neoantigen vaccination with vaccines designed according to the Jaime Leandro Foundation (JLF, v1 and v2) and CeGaT (Figure 1B, Table 1) protocols. The JLFv1 formulation contained nine peptides: five derived from single nucleotide variations (SNVs), and four derived from frameshift mutations (MYO15B, TECPR1.1, TECPR1.2, and MAP2). In addition to repeating all of JLFv1, JLFv2 added six additional targets, all derived from SNVs. Although untested in this work, the CeGaT vaccination series contained unique targets as well as targets overlapping with JLFv1 and JLFv2 (Figure 1C). Each vaccine formulation contained both HLA class I and class II predicted epitopes. Peptides contained within the JLFv1 vaccine were screened for pre- and post-vaccine IFNγ production following stimulation in culture; peptides found only in JLFv2 were only screened for post-vaccine reactivity.

**Figure 1:**
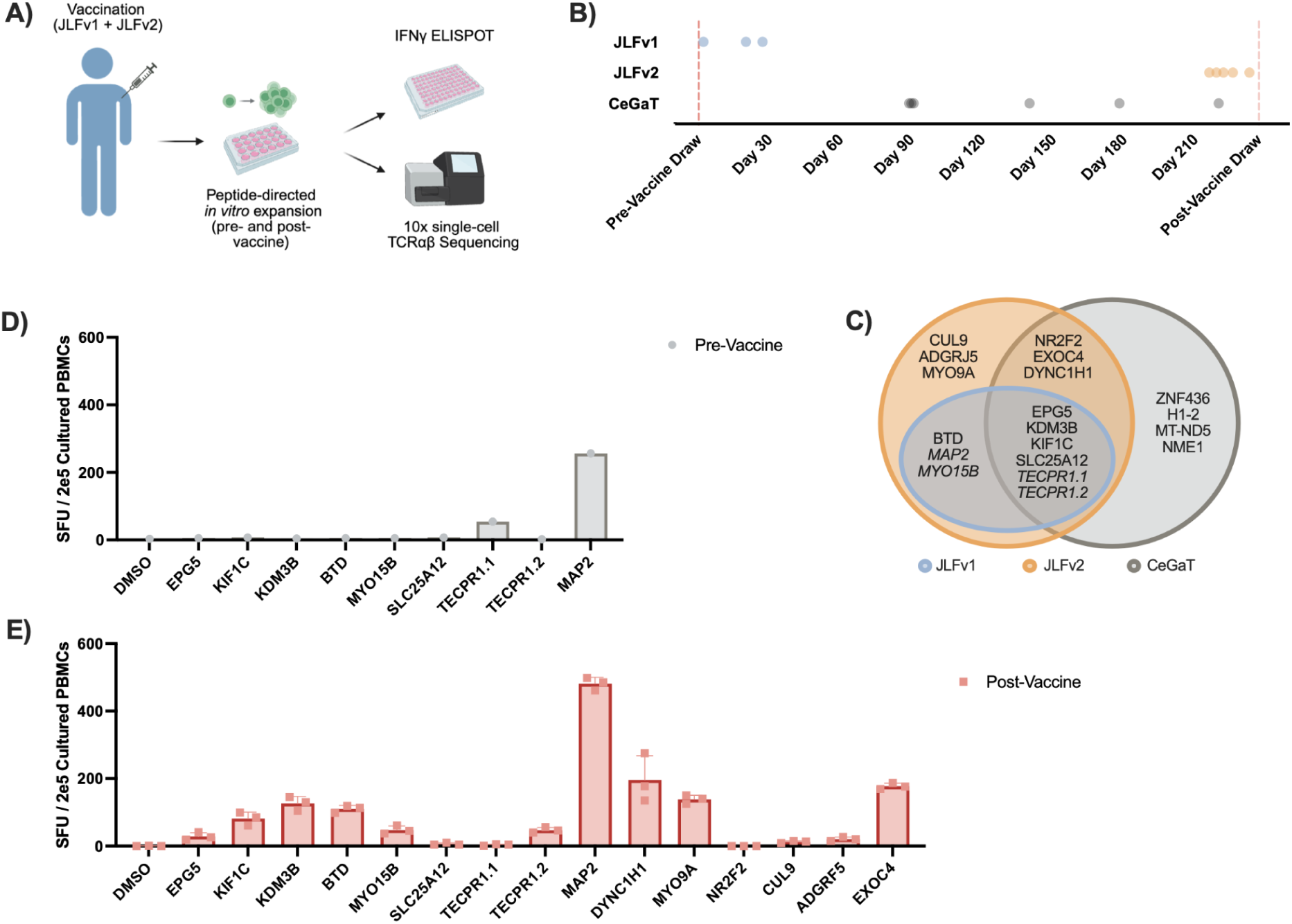
Personalized cancer vaccine design and post-vaccination immune monitoring. **A)** Schematic depicting workflow from personalized vaccination with neoantigen peptides to immune monitoring and TCR discovery via IFNγ ELISPOT and 10x single-cell TCRαβ sequencing. **B)** Timeline of vaccination dates for JLFv1, JLFv2, and CeGaT vaccine strategies. Sample collections used for immune monitoring analysis are indicated by dashed lines. **C)** Overlap of peptides between vaccine formulations. Frameshift mutations are represented in italics; all other mutations are SNV-derived. **D)** SFU from IFNγ ELISPOT of 2 × 10^5 cultured, peptide-pulsed PBMCs on day 11 of culture; pre- and **E)** post-vaccine timepoints. Post-vaccine co-cultures were submitted for 10x single-cell TCRαβ sequencing.

**Table 1:**
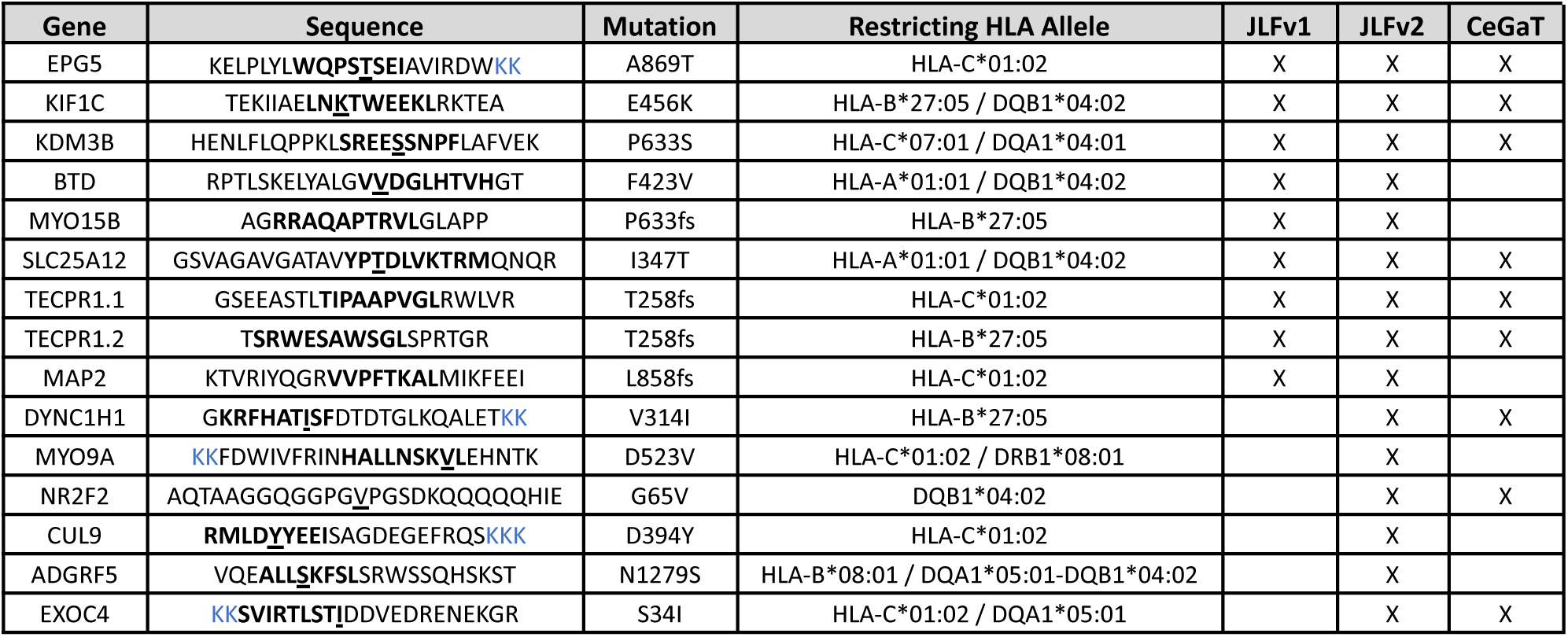
Neoantigen peptides administered as part of JLFv1 and JLFv2 vaccine formulations. Top predicted minimal epitopes (NetMHCpan v4.1b) are represented in bold; SNV mutations are underlined; additional modifications necessary for manufacturing are colored in blue. Shared vaccine targets with the CeGaT vaccine platform are also represented.

To investigate vaccine peptide reactivity, pre- and post-vaccine PBMC samples were viably thawed, stimulated with 10 ug/mL CSBio vaccine peptides, and cultured for 11 days, followed by an IFNγ ELISPOT with 2 × 10^5 cultured PBMCs / well. Of the peptides contained within the JLFv1 vaccine, TECPR1.1 and MAP2 elicited an IFNγ response in the pre-vaccine sample, although only the MAP2 peptide demonstrated strongly increased signal post-vaccination (Figure 1D, E, Supplemental Figure 1A, B). Additionally, peptides KIF1C, KDM3B, and BTD had moderately increased post-vaccine signal relative to the pre-vaccine timepoint. For peptides included only in the JLFv2 vaccine and so screened only for post-vaccine reactivity, IFNγ production was observed for DYNC1H1, MYO9A, and EXOC4 peptides (Figure 1E, Supplemental Figure 1B).

To maximize the probability of recovering an antigen-specific TCR, we sequenced all fifteen peptide-stimulated post-vaccine PBMC cultures using the 10x genomics single-cell TCR sequencing platform. To identify expanded T cell clones, 10x TCR single-cell sequencing results were analyzed using an in-house computational pipeline. After characterization based on CD4 or CD8 gene expression signatures, TCRs containing previously identified viral CDR3 regions were excluded from the dataset. We also observed a number of TCR clones, derived from both CD4^+^ and CD8^+^ T cells, that were found across multiple culture wells independent of stimulation; these TCRs were predicted to be non-specifically expanded and thus excluded from further analysis (Supplemental Figure 2A). Within these selection cutoffs, there was no observed relationship between IFNγ production and the frequency of the top 20 expanded TCR clonotypes for CD8^+^ T cell-derived TCRs, although CD4^+^ T cell-derived TCRs demonstrated a moderate correlation (Supplemental Figure 2B, C). Ultimately, filtering resulted in twenty-six TCRs from ten of fifteen peptide-stimulated pools to be generated and tested as TCR-T cells (Supplemental Figure 2D). Of note, there was no correlation between the number of TCR-Ts selected per peptide-pulsed stimulation and ELISPOT IFNγ signal.

### Antigen-specific TCRs are isolated from post-vaccination co-cultures

To further support the results of our immune monitoring IFNγ ELISPOT, we sought to determine the specificity of a set of clonally expanded TCRs isolated from the 11-day PBMC expansion cultures (Supplemental Table 1). Each of the twenty-six selected TCRs were expressed on primary CD8 T cells using sequential CRISPR-Cas9 knock-out of the endogenous TCR and retroviral transduction of the engineered TCR of interest (Figure 2A). Knock-out of the endogenous TCR (>99%) and expression of the virally transduced TCRs (25/26) were confirmed by flow cytometry (Supplemental Figure 3A, B).

**Figure 2:**
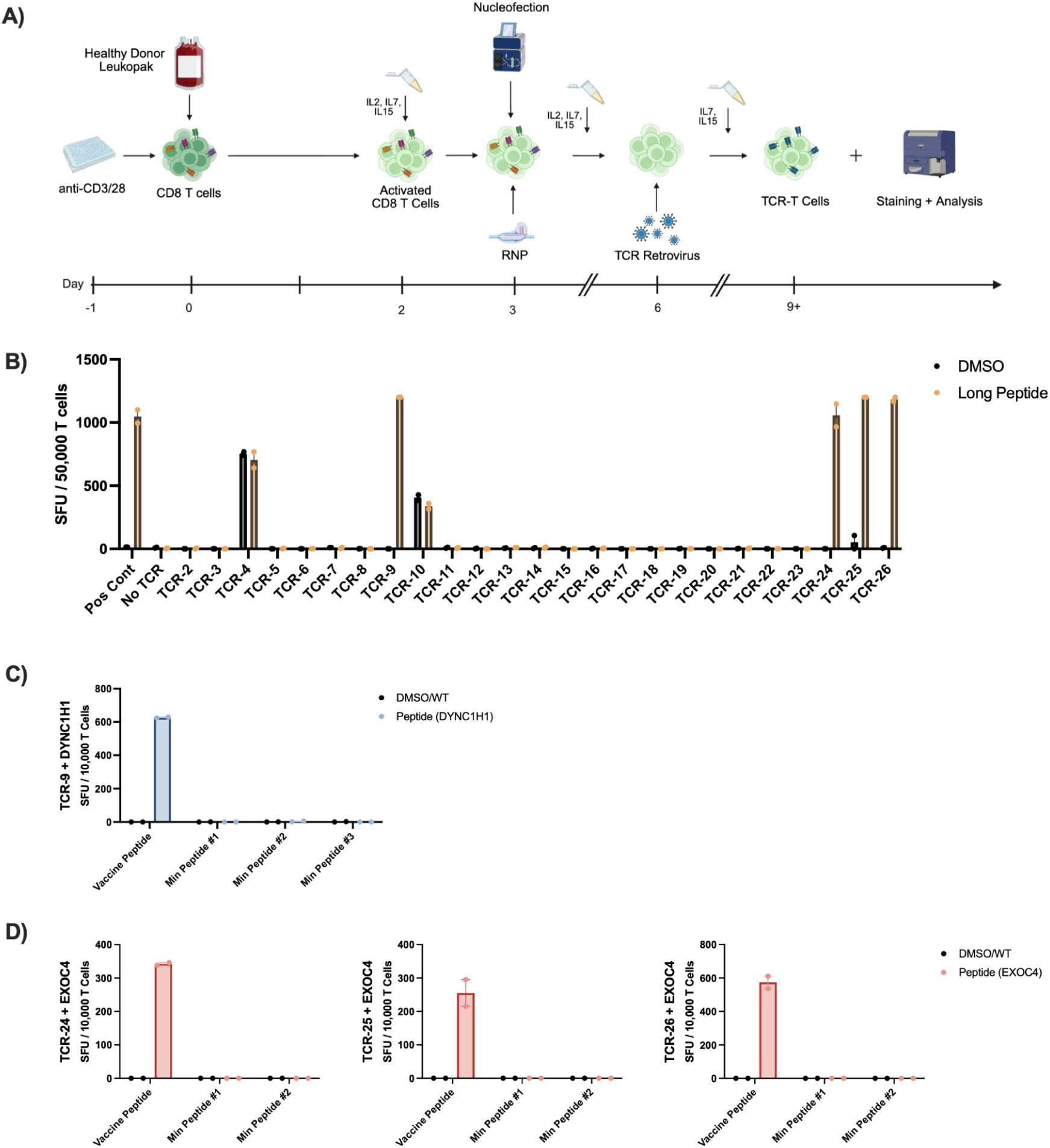
Selected TCR-T generation and testing. **A)** Schematic depicting TCR-T generation with CRISPR-Cas9 knock-out of endogenous TCR and retroviral transduction of novel TCR constructs. **B)** IFNγ ELISPOT of TCR-T and peptide-pulsed EBV-LCLs with vaccine long peptide or DMSO; SFU per 50,000 total T cells (n=2 technical replicates). **C)** IFNγ ELISPOT for positive TCR hit against vaccine long peptide DYNC1H1 and predicted minimal epitopes; SFU per 10,000 total T cells (n=2 technical replicates). **D)** IFNγ ELISPOT for positive TCR hits against vaccine long peptide EXOC4 and predicted minimal epitopes; SFU per 10,000 total T cells (n=2 technical replicates). Long peptides are compared to corresponding wild-type long peptides, minimal epitopes to DMSO

To evaluate the antigen-specificity of the generated TCR-T cells, we performed an IFNγ ELISPOT using autologous peptide-pulsed EBV-LCLs as antigen presenting cells (APCs). For 19 of 25 tested TCR-Ts, no increase in signal was observed in peptide-stimulated conditions compared to DMSO-pulsed controls. For two of the TCR-Ts, TCR-4D and TCR-10A, an IFNγ signal was detected in both the peptide-pulsed and DMSO conditions, likely indicating reactivity to an EBV-derived antigen presented by the EBV-LCLs. However, for four of the TCR-Ts (TCR-9, TCR-24, TCR-25, and TCR-26), a strong IFNγ signal was seen in only the peptide-stimulated wells (Figure 2B, Supplemental Figure 2C). TCR-9 was isolated from a co-culture stimulated with a long-peptide targeting an SNV within DYNC1H1; TCRs −24, −25, and −26 were all isolated from the co-culture stimulated with an EXOC4-derived long peptide, also containing an SNV. Of note, both co-cultures displayed moderate post-vaccine reactivity in the initial PBMC stimulation and ELISPOT, lower only than MAP2 stimulation (Figure 1E).

### PBMC culture-derived TCRs are specific to full-length vaccine peptides

In order to increase our understanding of TCR reactivity, we next set out to determine the minimal epitope for each of the four antigen-specific TCRs. To this end, we performed an additional series of IFNγ ELISPOTs, comparing predicted vaccine minimal epitopes to wild-type minimal epitope-pulsed EBV-LCLs (Table 2). For all four TCRs tested, the long vaccine peptide repeated a strong positive signal. However, none of the predicted minimal peptides resulted in IFNγ production. To verify that the minimal epitopes span the SNV mutation contained within the vaccine peptide, we tested each full-length wild-type peptide compared to the original vaccine peptide (Table 2). For each TCR, the original vaccine peptide repeated a strong positive signal by IFNγ ELISPOT, while the wild-type long peptide had little-to-no reactivity, suggesting a high degree of specificity to each of the SNV-derived neoepitopes (Figure 2C, 2D, Supplemental Figure 3D).

**Table 2:** Neoantigen peptides tested against TCR-9 (DYNC1H1) and TCRs −24, −25, and −26 (EXOC4). Predicted minimal epitopes are listed with their corresponding HLA proteoform and SNVs are underlined in both mutant and wild-type peptides.

|  | DYNC1H1 | EXOC4 |
| --- | --- | --- |
| Mutation | V314I | S34I |
| Vaccine Long Peptide | GKRFHAT<u>I</u>SFDTDTGLKQALET<u>KK</u> | <u>KK</u>SVIRTLST<u>I</u>DDVEDRENEKGR |
| Minimal Epitope #1 | <u>I</u>SFDTDTGL (HLA-C*01:02) | T<u>I</u>DDVEDRENE (HLA-B*27:05) |
| Minimal Epitope #2 | RFHAT<u>I</u>SF (HLA-B*27:05) | SVIRTLST<u>I</u> (HLA-B*27:05) |
| Minimal Epitope #3 | KRFHAT<u>I</u>SF (HLA-C*07:01) | n/a |
| Wild-Type Long Peptide | GKRFHAT<u>V</u>SFDTDTGLKQALET<u>KK</u> | <u>KK</u>SVIRTLST<u>S</u>DDVEDRENEKGR |

### Culture of PBMCs with vaccine peptides preferentially expands MHC-II restricted TCRs

To determine the minimal epitope contained within the vaccine long peptide, we ordered a comprehensive set of all possible 8-, 9-, 10-, and 11-mer peptides tiled across the SNV mutation site for both DYNC1H1 and EXOC4 (Genscript, custom). Due to the SNV’s proximity to the N-terminus, the DYNC1H1 set contained 32 peptides, compared to 38 peptides within the EXOC4 set (Figure 3A, Supplemental Figure 4A).

**Figure 3:**
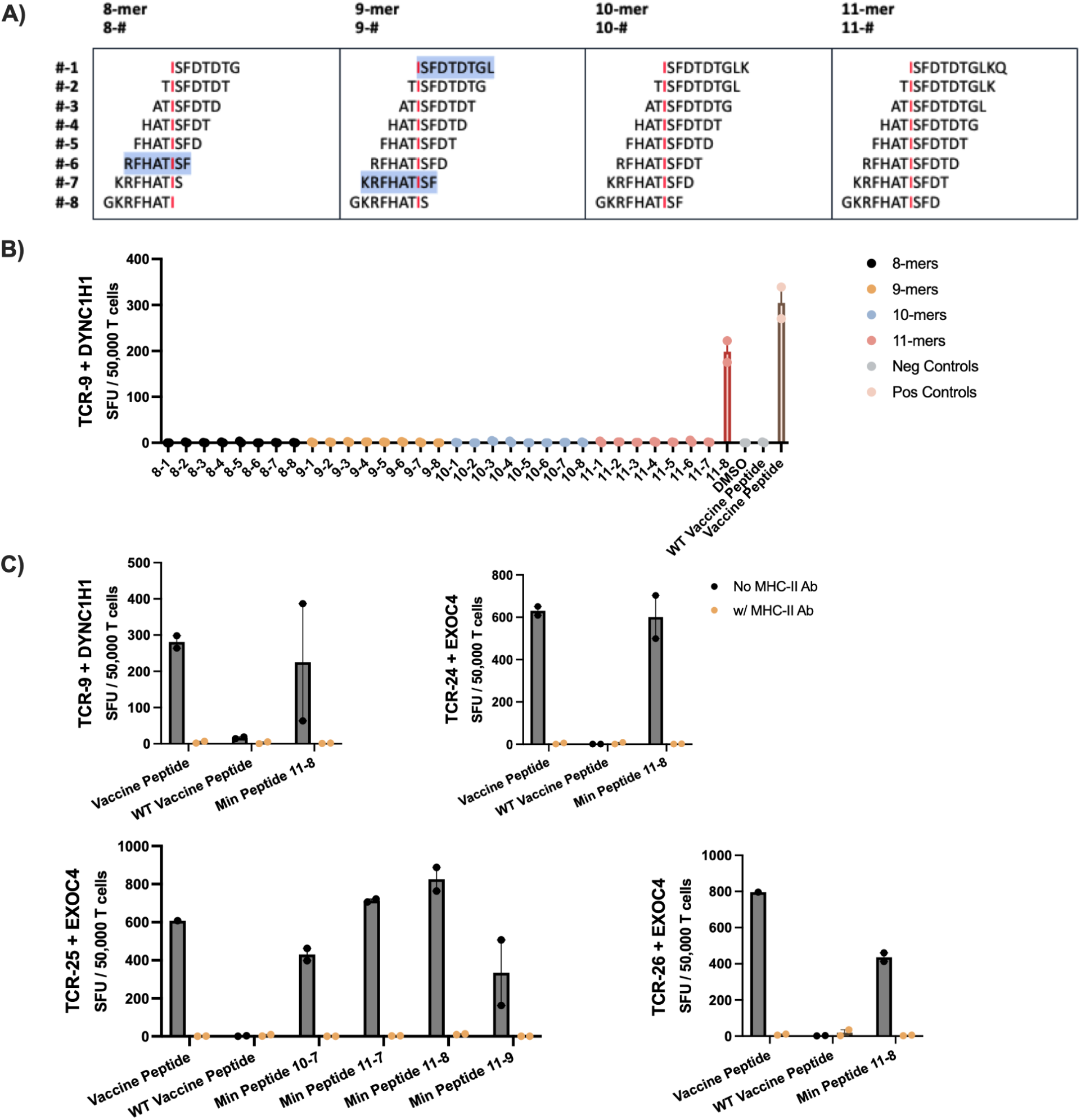
Determination of minimal epitopes for vaccine peptide reactive TCRs. **A)** Schematic depicting tiled class-I peptides across the DYNC1H1 mutation point (SNV, red). Previously tested minimal epitopes are highlighted in blue. Due to proximity of SNV to vaccine peptide end, only possible peptides are listed. **B)** IFNγ ELISPOT of TCR-9 against tiled DYNC1H1 class-I epitopes; SFU per 50,000 T cells (n=2 technical replicates). **C)** IFNγ ELISPOT of positive TCRs and corresponding minimal epitope hits, with or without a class-II blocking antibody; SFU per 50,000 T cells (n=2 technical replicates).

For all four TCRs, minimal reactivity was observed across all 8-, 9-, and 10-mers tested. However, within the possible 11-mers, each TCR produced an IFNγ signal above background for at least one epitope (Figure 3B, Supplemental Figure 4B, 5A-D). One TCR-T, −25, demonstrated reactivity across multiple adjacent epitopes (10-7, 11-7, 11-8, and 11-9). From these, we identified a ‘core’ shared sequence, IRTLSTIDD. We proceeded to compare this core epitope to predicted binding patterns across each of the patient’s class I HLA alleles (Supplemental Figure 6A). For four of the five alleles, no clear binding pattern was observable. The most likely binder, HLA-B*27:05, has a strong bias towards epitopes with an arginine (R) at position two. However, the core epitope possesses a negatively charged aspartic acid (D) residue at position 9, making it unlikely to bind HLA-B*27:05. To corroborate this qualitative analysis, we also ran the core epitope through NetMHCpan v4.1b. Although amongst the class I alleles the binding affinity (BA) was the “most” favorable for HLA-B*27:05 (BA = 27,570.03 nM), it would still be considered a non-binder by most metrics. By contrast, the core epitope was predicted as likely to bind to and be presented by an HLA class II allele, DRB1*08:01 (BA = 135.35 nM) (Supplemental Figure 4B). Of note, unlike the EXOC4 core motif, there are multiple class-I epitopes with strong predicted binding affinity contained within the DYNC1H1 reactive 11-8 peptide. However, the two most likely minimal epitopes, KRFHQTISF/HLA-C*07:01 (BA = 42.63 nM) and KRFHQTISF/HLA-B*27:05 (BA = 116.34 nM), were identified as non-reactive in both the initial predicted minimal epitope set and in the peptide-tiling screen (Figure 2C).

These data suggested that our four TCRs of interest may be specific to HLA class II epitopes rather than class I epitopes. To further evaluate this possibility, we performed another series of IFNγ ELISPOTs comparing the vaccine peptide, the corresponding wild-type peptide, and any minimal epitopes that previously elicited an IFNγ response, each incubated with and without an MHC-II blocking antibody (Figure 3C, Supplemental Figure 7A-D). For all peptide conditions across all TCRs, the class II blocking antibody completely abrogated the IFNγ response. To eliminate the possibility that a significant fraction of CD4^+^ T cells had expanded post-CD8^+^ T cell selection and were driving the IFNγ response, we performed intracellular cytokine staining (ICS) for CD8, CD4, and production of IFNγ, TNFα, and IL-2 following co-culture with the peptide-pulsed EBV-LCLs. We observed not only an expected absence of CD4^+^ T cells, but production of IFNγ, TNFα, and IL-2 by CD8^+^ T cells in response to long peptide stimulation (Supplemental Figure 7E). As such, we concluded that the IFNγ-producing TCRs were likely CD4^+^ T cell-derived and responding to epitopes presented by HLA-II. However, reactivity when expressed on CD8^+^ T cells suggests that the TCRs are co-receptor independent.

### Isolated CD8^+^ T cell co-culture yields antigen-specific, class-I restricted TCRs

Given our interest in discovering TCRs targeting HLA class I presented antigens, we next set out to better optimize our co-culture conditions to facilitate improved discovery. To this end, we repeated our 11-day stimulation experiment as described above, with two separate adjustments designed to minimize CD4^+^ T cell signals. First, we set up co-cultures exactly as described above, but with the addition of MHC-II blocking antibody, refreshed every three days with media and cytokines. Our second set of cultures included CD8^+^ T cells, isolated via negative selection, co-cultured with autologous dendritic cells at a 4:1 E:T ratio (Figure 4A). For this experiment, due to limiting cell numbers, peptides were pooled in sets of five, with three pools per co-culture condition (Supplemental Figure 8A). To capture all possible expanded TCRs, each pool was sent directly to 10x TCR sequencing and analyzed as described above; to filter out any remaining CD4^+^ T cells, additional stringency was applied to selection of CD8^+^ gene signatures. Expanded clones were compared between the CD8^+^ T cell + DC condition, the PBMCs + MHC-II blockade condition, as well as any minimally expanded clones from the first round of analysis that fell below our first selection cutoff (Supplemental Figure 8B).

**Figure 4:**
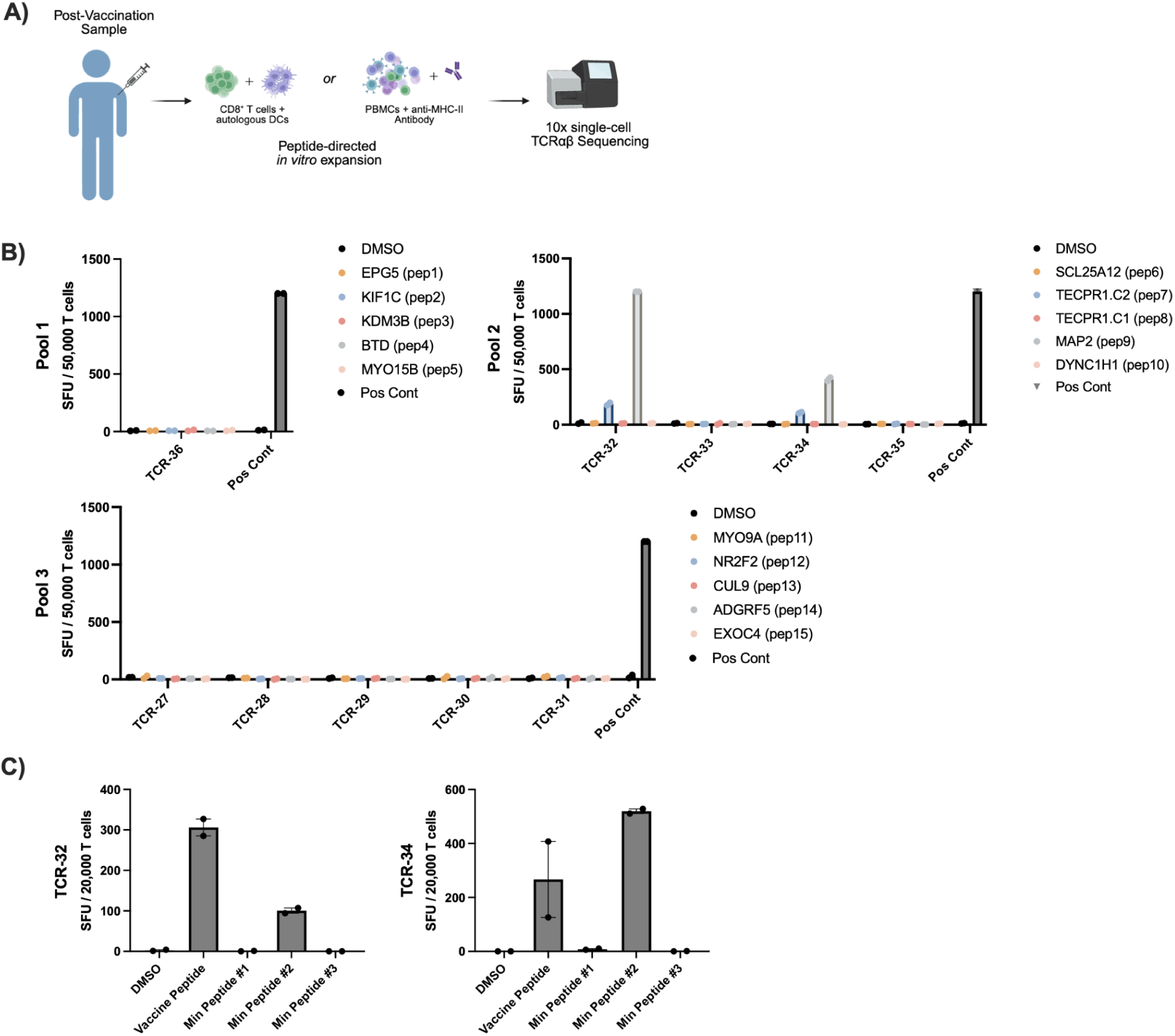
Improved TCR discovery conditions for isolation of vaccine peptide specific CD8-derived TCRs. A) Schematic depicting culture conditions for selectively expanding antigen-specific CD8^+^ T cells via pooled peptide stimulation. **B)** IFNγ ELISPOT of chosen TCRs against pooled peptides of origin; SFU per 50,000 T cells (n=2 technical replicates). **C)** For TCRs reactive to long peptide stimulation, IFNγ ELISPOT against corresponding predicted minimal epitopes; SFU per 50,000 T cells (n=2 technical replicates).

This second round of stimulation yielded 10 new TCRs for downstream TCR-T generation and testing. Each of the three pools of stimulation was represented (Pool 1 = 1 TCR; Pool 2 = 4 TCRs; Pool 3 = 5 TCRs). Of these ten TCRs, retrospective analysis found that only four were present in the original 15-peptide whole PBMC screen. Amongst these four, three were previously eliminated from further testing due to detection in two or more individually stimulated cultures (TCRs −27, −31, and −33). The remaining overlapping TCR was assessed as below the limit of selection in the initial screen, present as just 0.01% of the TCR repertoire (Supplemental Figure 8B). To deconvolute antigen specificity, TCR-Ts were screened via IFNγ ELISPOT against each of the five peptides used in the original stimulation pool. Although no TCR-Ts demonstrated reactivity within Pool 1 or Pool 3, two TCR-Ts from Pool 2 reacted to the frameshift-containing MAP2 vaccine peptide (Figure 4B, Supplemental Figure 9A). Of note, upon further testing with three predicted minimal epitopes, both TCRs demonstrated strong reactivity to minimal peptide #2 (Figure 4C, Supplemental Figure 9B).

## Discussion

This study demonstrates that antigen-specific T cell populations, bolstered by neoantigen peptide vaccines, are targetable as a source of T cell receptors for personalized cellular therapies. In this single patient study, a locally reccurent osteosarcoma patient received three courses of personalized neoantigen-targeting synthetic long peptide vaccines, administered externally by the Jamie Leandro Foundation (JLFv1 and JLFv2) and CeGaT (CancerNeo). We observed increased magnitude of immune responses to both SNV-derived and frameshift mutations following neoantigen peptide vaccination for seven of ten neoantigens tested by IFNγ ELISPOT (Figure 1D, 1E). Of note, the strongest IFNγ response observed was against the MAP2 peptide, which was derived from a frameshift mutation. A strong response was observed against this peptide in the pre-vaccine sample, suggesting that the original mutation was sufficiently immunogenic to elicit a detectable *in vivo* immune response. Following vaccination, we observed that the response was further elevated, suggesting that the peptide vaccination successfully bolstered the existing immune response. This finding supports the hypothesis that frameshift mutations can be particularly immunogenic, as they elicit an entirely novel protein sequence with little homology to self proteins. The remaining six boosted peptides, for which a signal was not detected in the pre-vaccine sample but which was detected in the post-vaccine sample, contained two additional frameshift derived mutations (MYO15B, TECPR1.2) and four SNV derived mutations. Although constraints on sample availability limited our ability to test pre-vaccine samples on the full JLFv2 vaccine formulation, we detected strongly positive responses (SFU > 50) for three additional peptides post-vaccination. Taken together, these results further support the immunogenicity of the peptide vaccine formulation in bolstering the *in vivo* immune response such that it becomes observable within conventional limits of detection.

The approaches used here build upon a number of established protocols^2,41,42,48^. In using total PBMCs for a direct *ex vivo* ELISPOT, particularly when stimulating cultures with long (>15 AA) peptides, IFNγ responses may be driven by CD4^+^ T cells, CD8^+^ T cells, or a combination thereof. It is well established that only the strongest immune responses are detectable directly *ex vivo*^49,50^; as such, many groups have adopted an *in vitro* stimulation (IVS) approach. Options for IVS cultures include total PBMCs, isolated T cells with autologous APCs (dendritic cells or EBV-LCLs), or isolated T cells with engineered or artificial APCs. Use of total PBMCs is a standard approach for general immune monitoring, as it maximizes all available cells and is sufficient to detect both pre- and post-vaccine responses (Figure 1D, E). In contrast, use of an isolated T cell compartment with matched APCs allows for determination of signal from a specific T cell population. However, these approaches have significant limitations. While autologous dendritic cells have long been regarded as the gold standard of antigen presentation, they are a non-renewable resource and require significant sample input to generate at scale. Alternatively, autologous EBV-LCLs are an entirely renewable tool, requiring minimal sample input to generate. However, extended use of EBV-LCLs as APC risks expansion of T cell populations targeting EBV rather than the antigen(s) of interest. All of these issues may be mitigated by use of an artificial APC, such as an engineered K562 cell line. However, this approach requires generation of a cell line expressing each HLA of interest, quickly becoming cost prohibitive for most studies involving personalized therapies.

Here, we elected to maximize sample efficiency by performing an 11-day *in vitro* stimulation of total PBMCs for pre- and post-vaccine samples. Although this approach was sufficient for detection of total T cell responses for immune monitoring, we encountered challenges in our secondary aim: to identify and characterize CD8^+^ T cell-derived TCRs. Bioinformatics analysis of post-vaccine PBMC cultures selected twenty-six expanded TCRs for downstream testing, chosen by criteria including CD8^+^ expression, culture specificity, and removal of known viral CDR3 sequences (Supplemental Figure 2A, D). Notably, this selection process yielded only four vaccine antigen-specific TCRs out of twenty-five validated constructs, suggesting that our selection based primarily on clonal expansion remains suboptimal. Even within those four TCRs, further characterization revealed that each was CD4^+^ T cell-derived, and as such, restricted to antigens presented on MHC class-II. Although outside the scope of this work and thus not further investigated within this study, the clinical efficacy and utility of co-receptor independent CD4^+^ T cell-derived TCRs remains to be determined.

Given our interest in identifying TCRs capable of targeting HLA class I presented antigens, we set out to determine a set of culture conditions that could be used to preferentially stimulate CD8^+^ T cells within the constraints of limited starting cell numbers. First, we generated autologous dendritic cells to be cultured with isolated CD8^+^ T cells. For a second set of culture conditions, we again used whole PBMCs; however, we aimed to support preferential CD8^+^ T cell stimulation and expansion by inclusion of MHC-II blocking antibody. This second round of TCR selection yielded ten novel TCRs for testing and validation. Within the set, nine were selected for expansion from the DC:CD8^+^ T cell co-cultures; only one was selected from the PBMC + anti-MHC-II cultures. The ten novel TCR-Ts were generated and screened via IFNγ ELISPOT, with each TCR tested against each of the five peptides originally used to stimulate its pool of origin. Two of the ten TCRs, −32 and −34, tested were reactive to vaccine long peptide #9, containing a MAP2 derived frameshift mutation. Of note, neither of these TCRs was detected at any frequency in the initially expanded post-vaccine PBMC sample, and both were isolated from the DC:CD8^+^ T cell co-cultures (Supplemental Figure 8A, B). Further screening of the TCRs against predicted minimal epitopes showed that both TCRs were reactive to minimal epitope GRVVPFTKA, expressed on HLA-B*27:05. Although not investigated further here, we observed that TCR-34 demonstrated higher reactivity to the minimal epitope than to the vaccine long peptide, while the opposite was true for TCR-32. We predict this is an indicator that TCR-32’s true minimal epitope likely overlaps imperfectly with our predicted minimal epitope, while the vaccine long peptide contains the full, true minimal epitope, leading to an increased IFNγ signal.

Our study is limited in important ways. Primarily, this work represents a single case in which neoantigen peptide vaccination elicited T cell responses of sufficient magnitude to be detectable via IFNγ ELISPOT and support discovery of a tumor antigen-specific TCR. Experiments were limited by the availability of primary patient samples, particularly from the pre-vaccine timepoint. Further studies would be necessary to understand how both the immunogenicity of the chosen vaccine peptides and the magnitude of the T cell response generalize to TCR discovery in a larger patient population. Although outside the scope of this work, each of these concepts should also be examined in line with the selection process for vaccine peptides as well as vaccine administration with respect to disease status. While many of these limitations are inherent to personalized therapies, particularly at the developmental stage, they do not diminish their importance for select patients.

Personalized antigen-directed immunotherapies offer a path forward for many patients for whom standard-of-care therapeutics have failed. A significant advantage of such approaches is their theoretical capability to capitalize on an uncapped range of HLA haplotypes and mutations. However, all forms of antigen-directed immunotherapies rely deeply on antigen selection, and thus remain limited in potential by the narrow body of literature available to train binding and presentation prediction models, which is heavily biased towards commonly expressed HLA alleles, leading to lower confidence predictions for uncommon HLAs^51,52^.

Careful antigen selection is of particular importance for neoantigen vaccinations, for which little-to-no experimental validation of target immunogenicity is typically performed prior to dosing; rather, immunogenicity is often assessed post-hoc via immune monitoring experiments, as demonstrated here and by others^11,41,42^. In this work, we have provided evidence in support of both immunogenicity of cancer peptide vaccines as well as the JLF antigen selection algorithms, demonstrating increased immune responses in 80% of vaccine peptides screened pre- and post-vaccination. Notably, we demonstrated immune responses ranging across predictions for each of the five HLA alleles represented in this patient. This encompasses four common HLA alleles, defined as the top 10 most frequently occurring worldwide by gene, as well as responses to peptides containing mutations predicted to be expressed on a comparatively less common HLA allele, HLA-B*27:05. This dispersed hit frequency across all HLA alleles highlights an advantage of neoantigen vaccines: the ability to include a large range of predicted target antigens spread across a patients’ unique set of HLA proteoforms. Recent trials have included up to 20 individual targets for peptide vaccines^41,53^, and up to 34 targets for mRNA vaccine platforms^11,12,16,54^. In addition to decreasing the impact of a single, non-immunogenic predicted antigen, inclusion of a broader range of antigen targets also mitigates the risks posed by loss of heterozygosity affecting a single HLA allele or an entire phased haplotype.

By comparison, TCR-T-based therapies have historically been limited to a single TCR targeting one peptide-HLA complex, although more recent studies have demonstrated safety with products containing up to three unique TCR-Ts^10^. This is due in part to the significant challenges and expenses associated with personalized TCR discovery, as well as a lack of applicable ‘off-the-shelf’ TCR-T therapies for any given patient. Despite becoming increasingly available over the last decade, TCR-T therapies targeting public neoantigens are heavily restricted in use, not only by expression of a specific mutation in any given tumor, but also by the requirement that a patient have an exact match for the corresponding HLA proteoform. Beyond challenges in HLA matching, public neoantigens for semi-personalized use are most commonly identified within driver mutations. Some of the most frequently occurring public neoantigens are derived from driver mutations occurring within *BRAF*, *KRAS*, PIK3CA, and *TP53*^55,56^. A number of TCRs specific to these neoantigens across a range of HLAs have been characterized^27,57,58^, with some demonstrating clinical efficacy when deployed as TCR-Ts^7,18,59^. While the availability of even a small number of semi-personalized cellular therapies represents tremendous progress, it also highlights the need to continue developing personalized antigen-directed cellular therapies. As demonstrated here, discovery of TCRs for a single patient is able to extend beyond commonly shared targets into individual mutations, regardless of shared population prevalence, while also targeting mutation types beyond SNVs.

In conclusion, our study offers insight into means of discovery of highly personalized TCRs targeting an immunogenic frameshift mutation, with potential for use as a downstream cellular therapy. We also offer insights into a combinatorial approach of a personalized cancer vaccine to prime the immune response prior to isolation of TCRs, as well as highlighting differential approaches for isolation of CD8^+^-specific T cell responses under sample-constrained conditions. Although further testing is needed to determine clinical applicability and *in vivo* functionality, this work supports the future development of personalized TCR-T therapeutics following therapeutic vaccination.

## Declarations

### Availability of data and materials

Sequencing data and other raw data will be made available upon request to the corresponding author.

### Authors’ contributions (CRediT roles)

SNB: Conceptualization, Formal Analysis, Investigation, Methodology, Visualization, Writing - original draft, Writing - review & editing

JA: Investigation, Methodology, Writing - review & editing

MS: Investigation, Methodology

JG: Methodology KMK: Investigation

AS: Project Administration, Resources

WBI: Project Administration, Resources

WH: Project Administration, Resources, Funding Acquisition

MT: Project Administration, Resources, Funding Acquisition

AR: Conceptualization, Data Curation, Formal Analysis, Funding Acquisition, Methodology, Software, Supervision, Visualization, Writing - review & editing

BGV: Conceptualization, Funding Acquisition, Project Administration, Resources, Supervision, Writing - review & editing

## Acknowledgements

We would like to acknowledge the work and contributions of several core facilities at UNC, including the Immune Monitoring and Genomics Facility (IMGF), the High Throughput Sequencing Facility (HTSF), and the Flow Cytometry Core Facility. We also thank (M.S.) for their assistance in reviewing the manuscript.

## List of Abbreviations

TCR: T Cell Receptor
TCR-T: T Cell Receptor-engineered T cell therapy
PBMC: Peripheral Blood Mononuclear Cell
ELISPOT: Enzyme-Linked ImmunoSpot
MHC: Major Histocompability Complex
HLA: Human Leukocyte Antigen
PCV: Personalized Cancer Vaccine
ETC: Endogenous T Cell therapy
TIL: Tumor-Infiltrating Lymphocyte
SNV: Single Nucleotide Variant
SFU: Spot Forming Unit
UMI: Unique Molecular Identifier
GEX: Gene Expression data

---

**Supplemental Figure 1:**
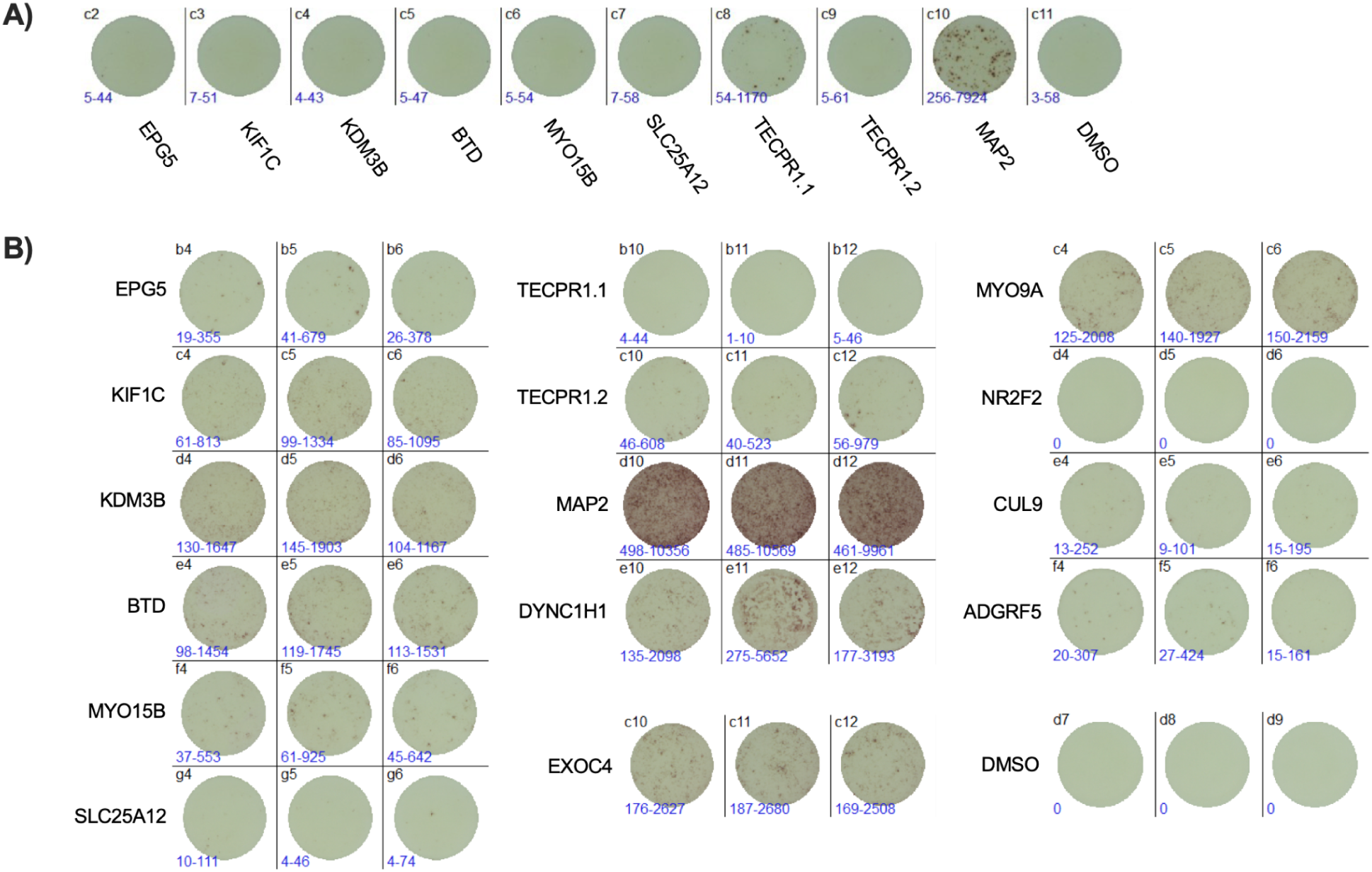
Immune monitoring IFNγ ELISPOT results. **A)** Raw images of IFNγ ELISPOT wells from 2e5 cultured, peptide-pulsed PBMCs on Day 11, pre- and **B)** post-vaccine timepoints.

**Supplemental Figure 2:**
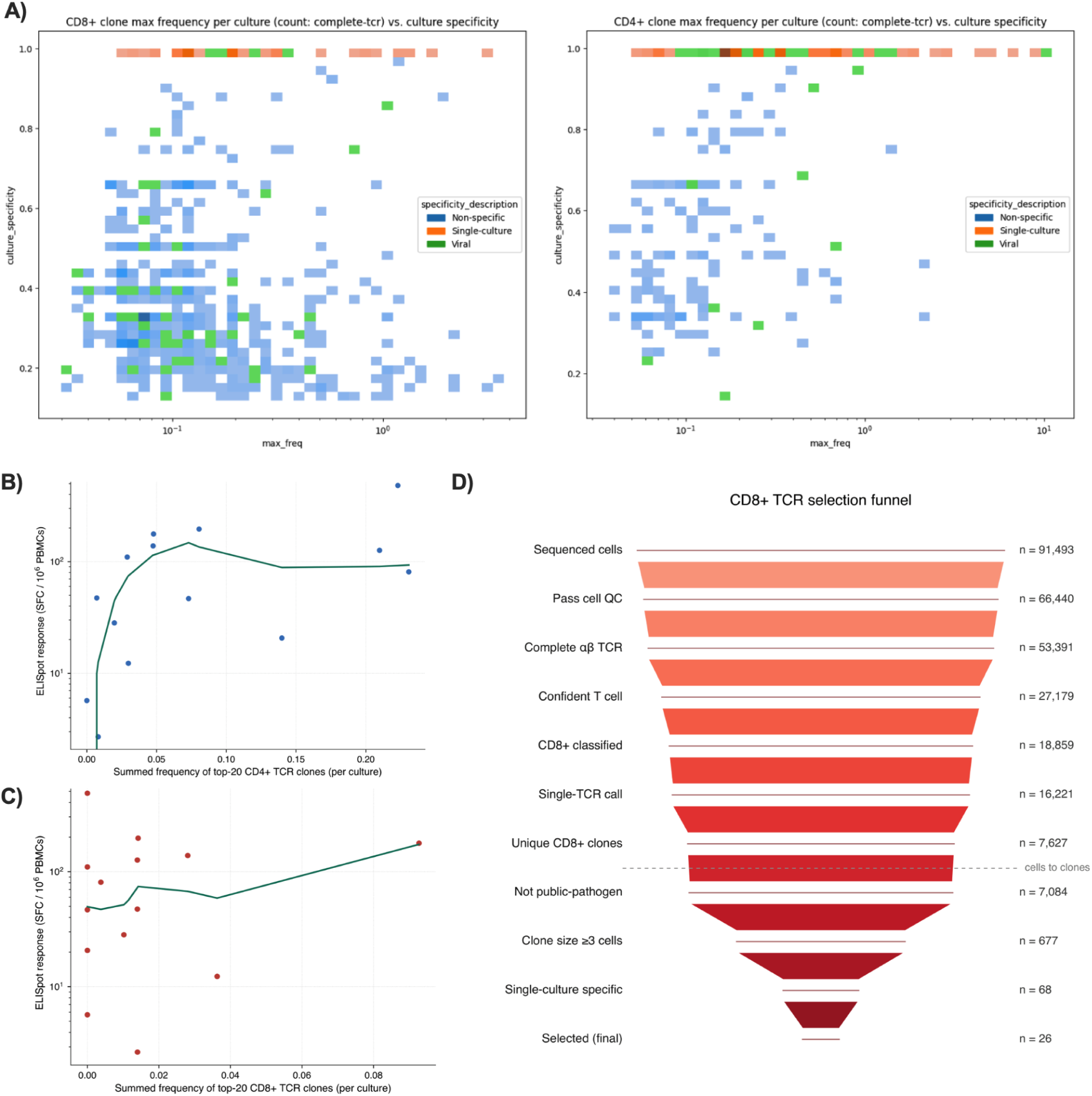
Analysis and selection of TCRs from vaccine peptide-pulsed PBMC cultures. **A)** Logistic model of frequency vs occurrence of CDR3β sequences from 10x single-cell TCRαβ sequencing data for CD4^+^ and CD8^+^ T cells. CDR3β sequences previously characterized as virally associated are depicted in green; sequences found in multiple peptide-stimulated cultures are indicated in blue. **B)** ELISPOT response vs per-culture summed clonal expansion of top-20 CD4^+^ TCR clones and **C)** CD8^+^ TCR clones. **D)** Selection funnel depicting steps from total sequenced cells to n=26 TCR sequences selected for downstream analysis.

**Supplemental Figure 3:**
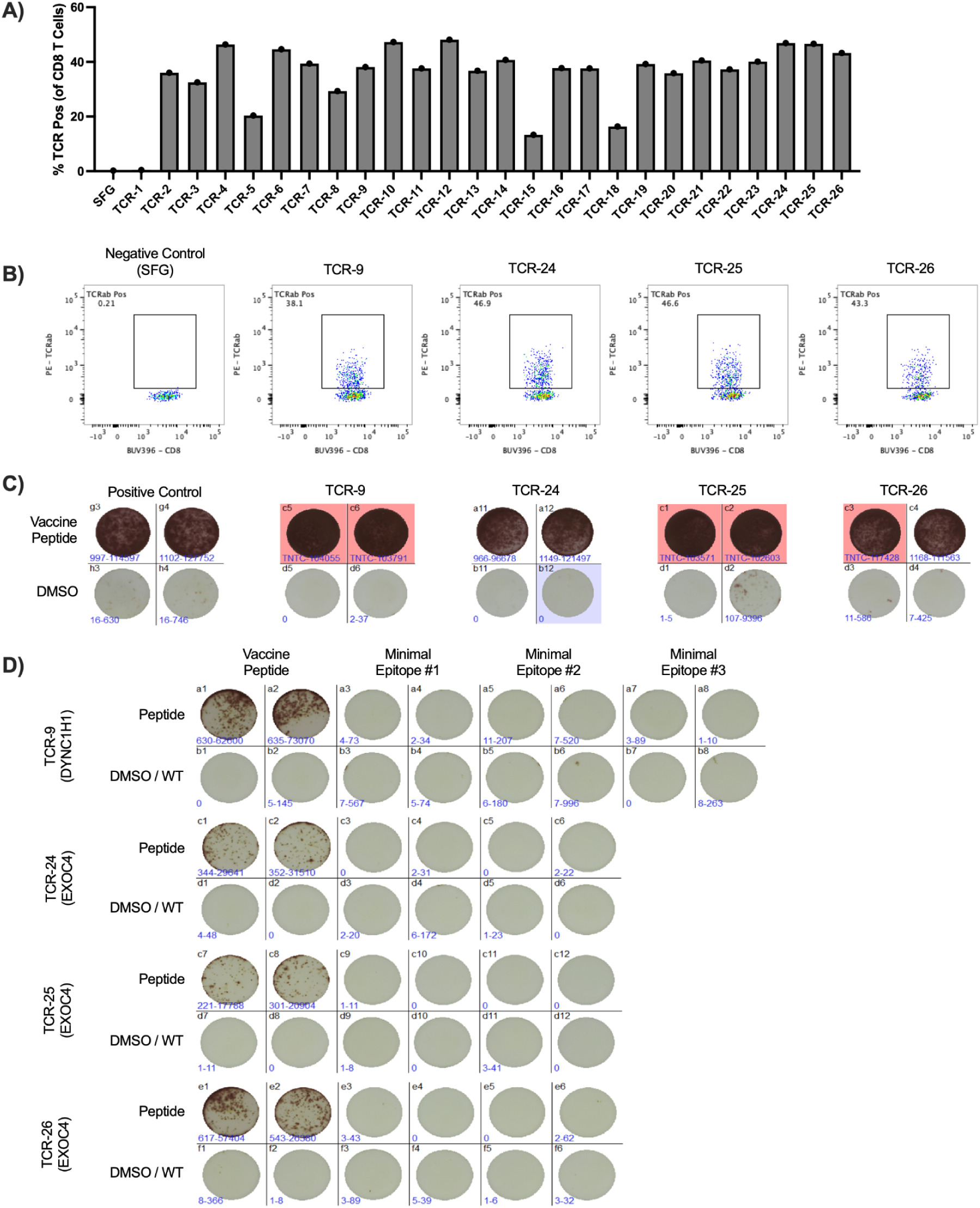
Supporting data for selected TCR-T generation and testing. **A)** Surface expression of TCRa/b as a percentage of live, CD8^+^ cells. **B)** Representative flow plots of CD8 x TCRa/b staining for TCRs −9, −24, −25, and −26. Briefly, cells were gated as lymphocytes, singlets, live, CD8^+^, and presented as percent TCRαβ positive of live, CD8^+^ T cells. **C)** Representative images of IFNγ ELISPOT wells for TCRs −9, −24, −25, and −26, as well as an internal positive control TCR stimulated with vaccine long peptides or DMSO; SFU per 50,000 total T cells (n=2 technical replicates). **D)** Raw images of IFNγ ELISPOT wells stimulated with vaccine peptides, predicted minimal epitopes, and wild-type minimal epitopes for TCRs −9, −24, −25, and −26; SFG per 10,000 total T cells (n=2 technical replicates).

**Supplemental Figure 4:**
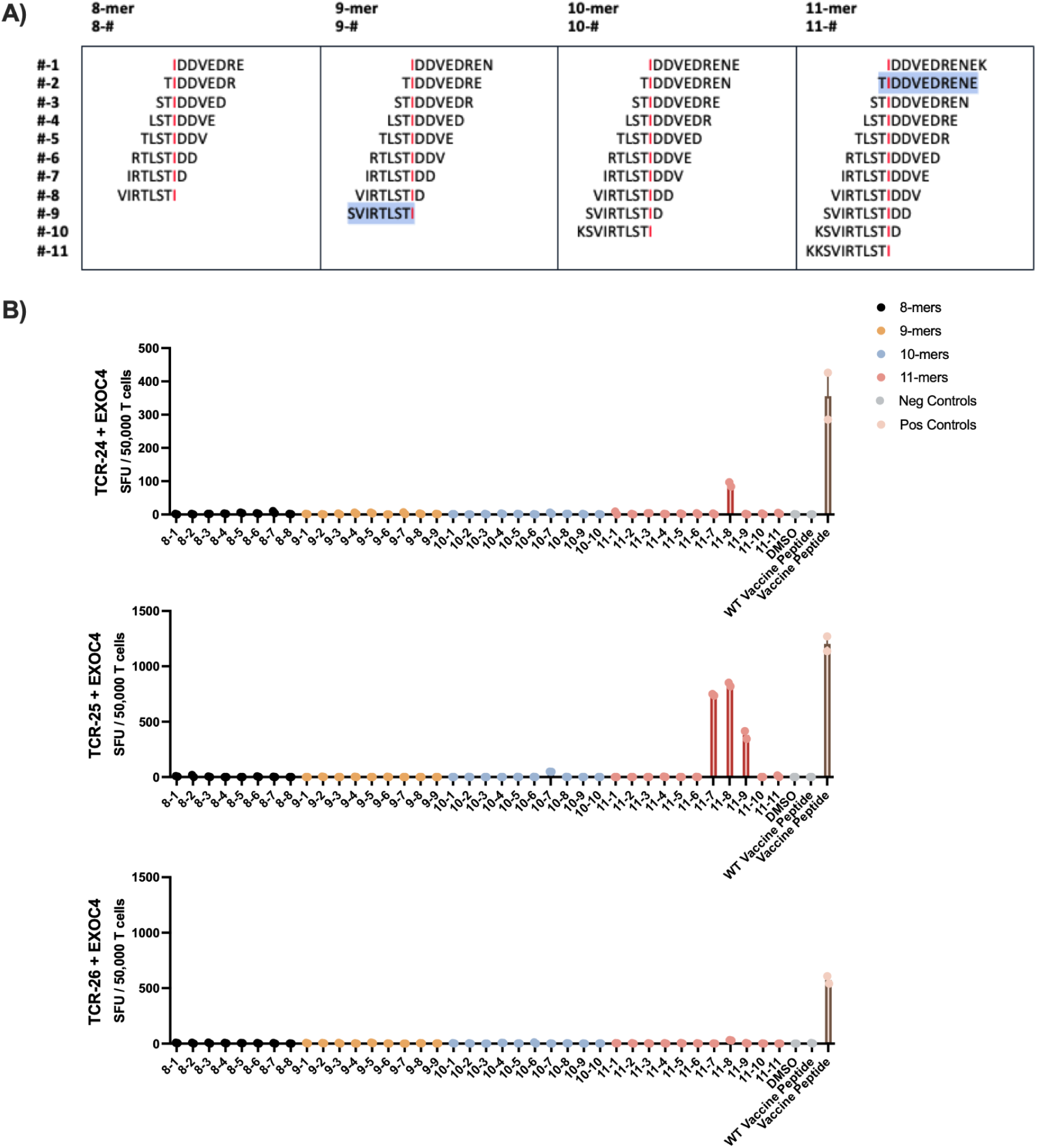
Determination of minimal epitopes for EXOC4 reactive TCRs. **A)** Schematic depicting tiled class-I peptides across the EXOC4 mutation point (SNV, red). Previously tested minimal epitopes are highlighted in blue. **B)** IFNy ELISPOT of TCRs −24, −25, and −26 against tiled EXOC4 class-I epitopes; SFU per 50,000 T cells (n=2 technical replicates).

**Supplemental Figure 5:**
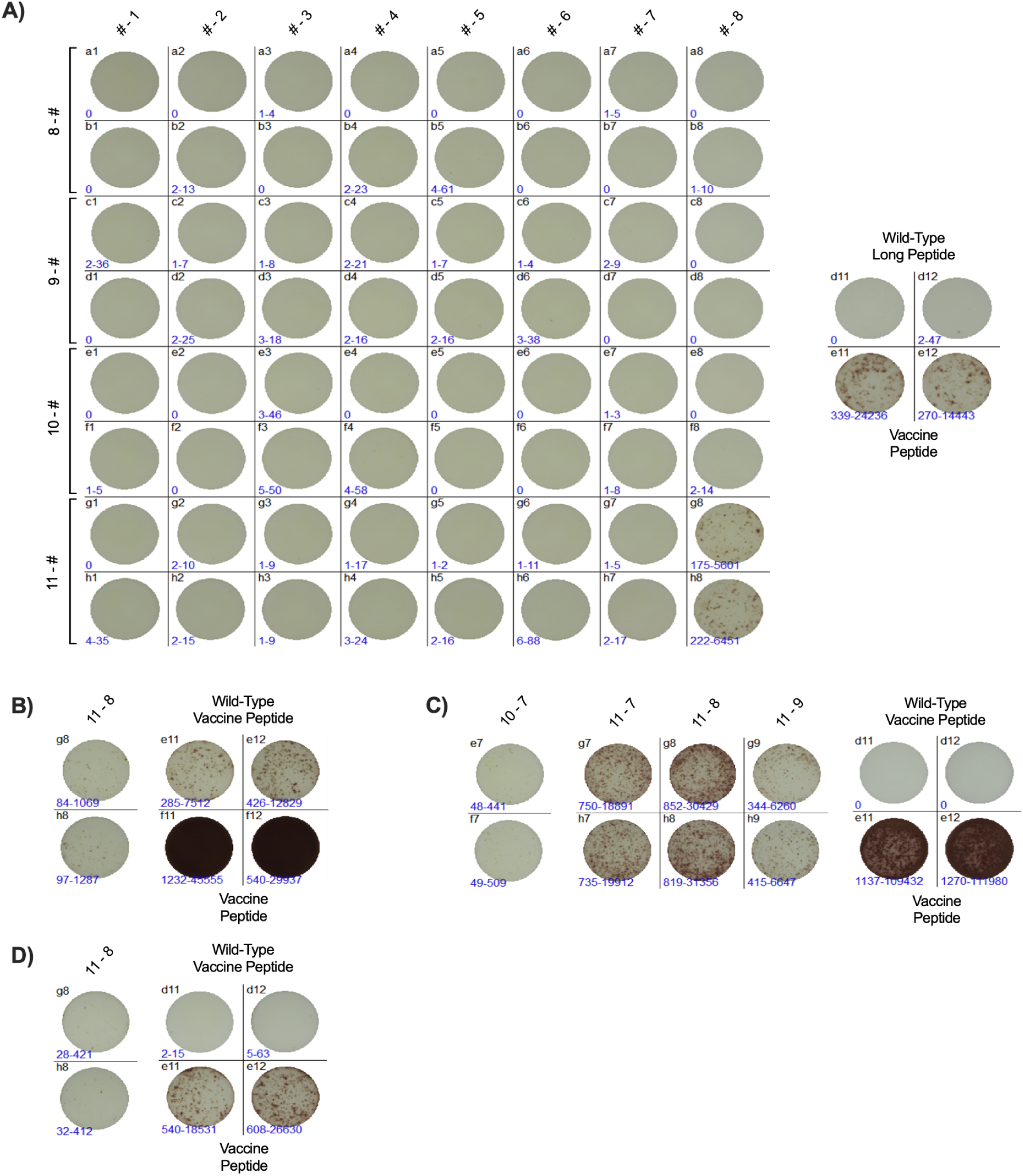
Supporting raw ELISPOT data for peptide tiling to assess minimal epitopes. **A)** Raw images of peptide tiling across peptide DYNC1H1 with TCR-9, assessed by IFNγ ELISPOT; SFU per 50,000 T cells (n=2 technical replicates). **B)** Representative raw images of positive IFNγ ELISPOT wells of TCR-24, **C)** TCR-25, and **D)** TCR-26; SFU per 50,000 T cells (n=2 technical replicates).

**Supplemental Figure 6:**
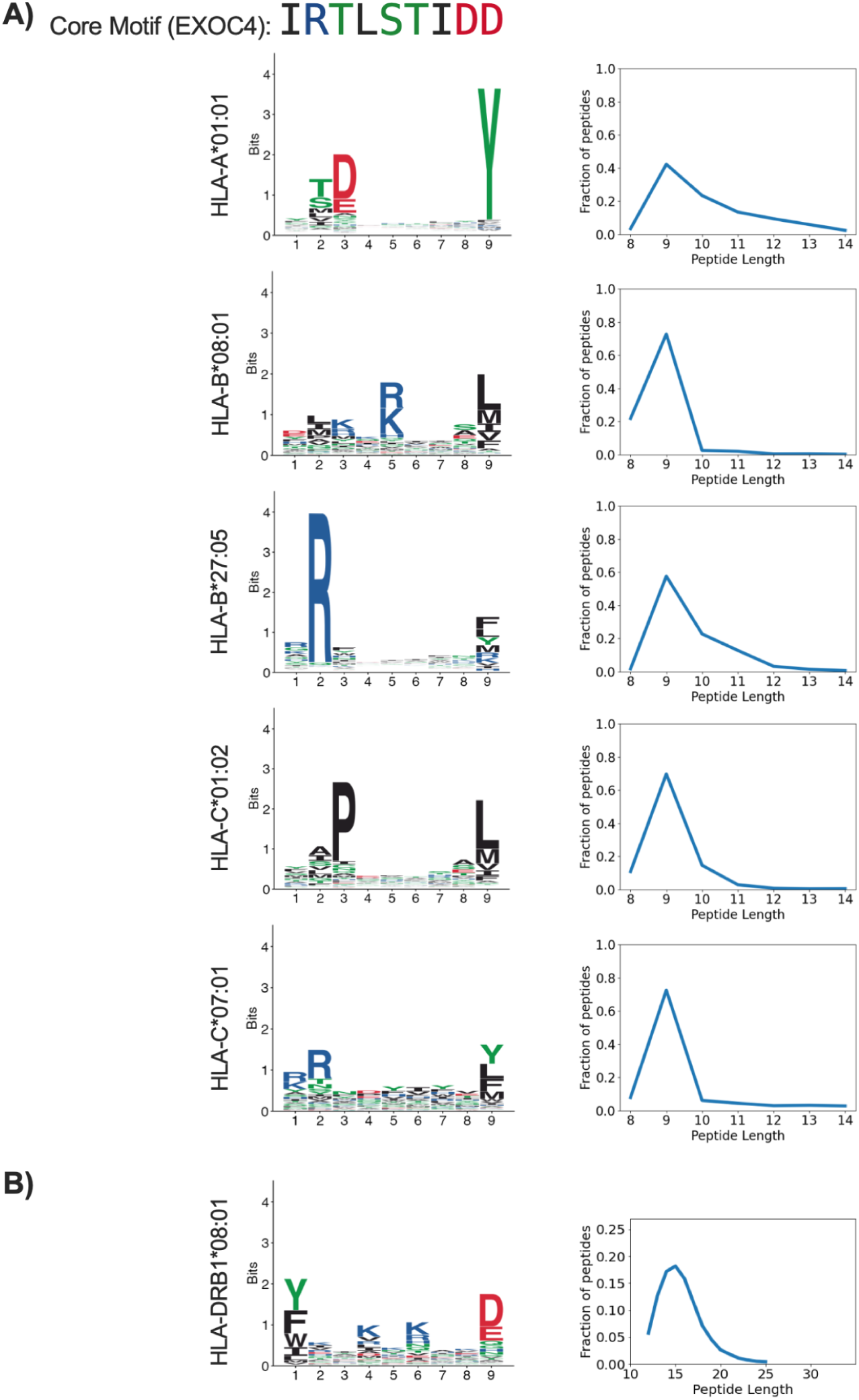
Assessment of EXOC4 core epitope HLA binding. **A)** Identified EXOC4 core epitope and predicted MHC binding motifs for possible class I alleles. **B)** Predicted MHC binding motif for class II allele DRB1*08:01. All MHC binding motifs sourced from The MHC Motif Atlas^60^.

**Supplemental Figure 7:**
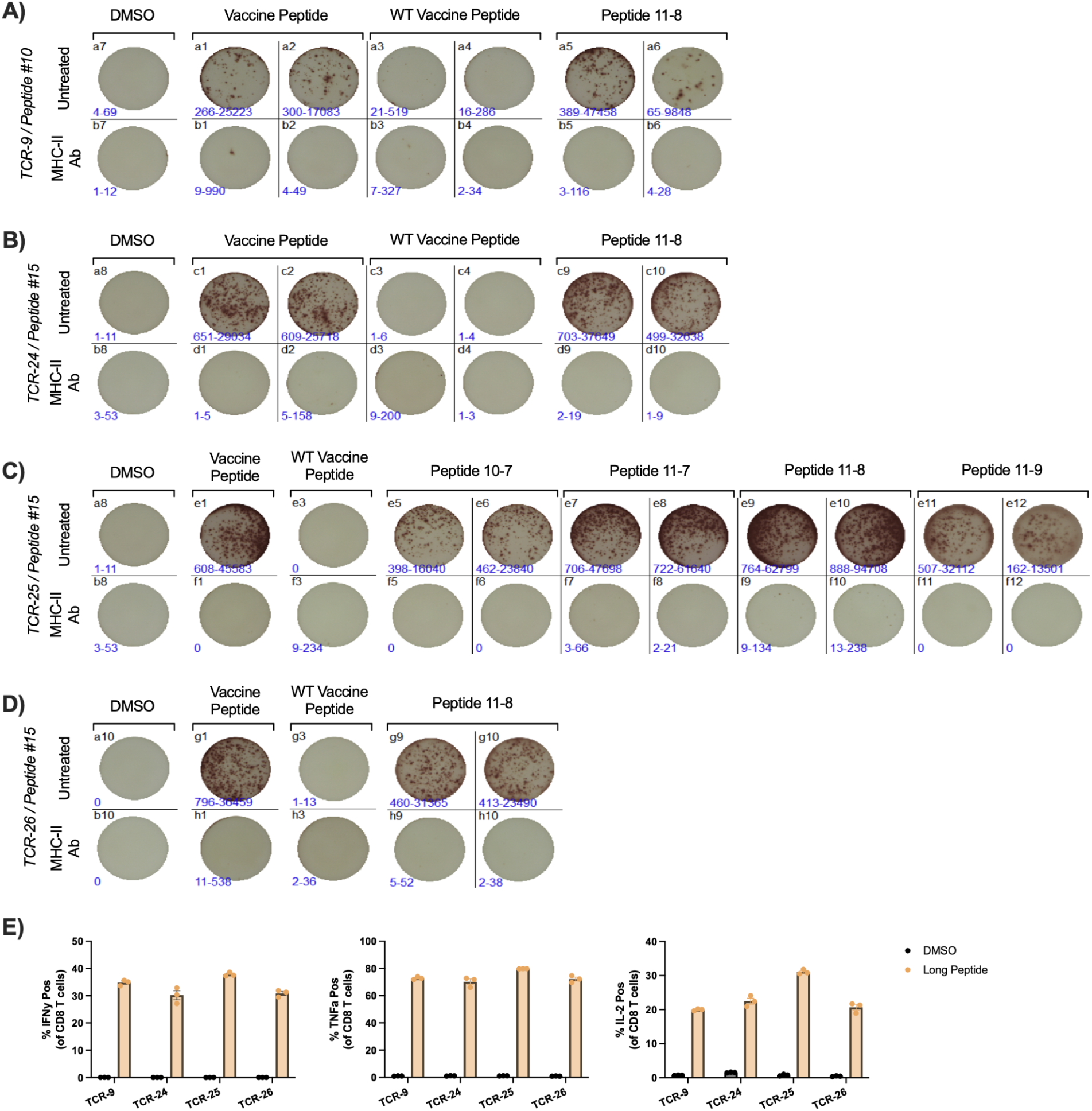
Supporting data for classification of TCRs as MHC class-II restricted. Representative raw images of IFNγ ELISPOT with and without class-II blocking antibody (SFU per 50,000 T cells, n=2 technical replicates) for **A)** TCR-9, **B)** TCR-24, **C)** TCR-25, and **D)** TCR-26. Note: TCR-25 and −26 have single replicates for long peptide controls. **E)** Intracellular cytokine staining of CD8^+^ TCRT cells for IFNγ, TNFα, and IL-2. Cells were gated as lymphocytes, singlets, live, CD8^+^, and presented as percent cytokine positive of live, CD8^+^ T cells.

**Supplemental Figure 8:**
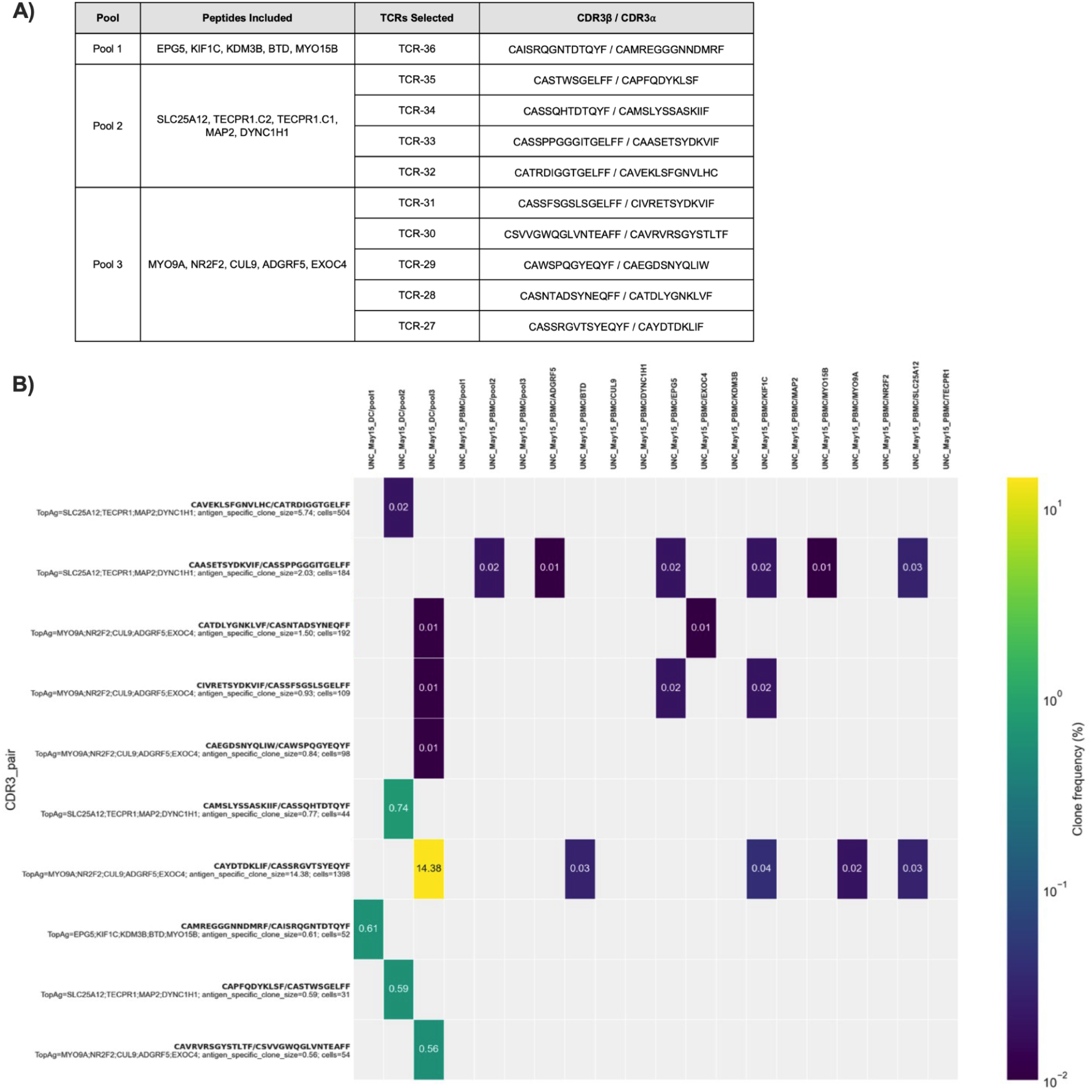
TCR Selection from optimized TCR discovery conditions. **A)** Table depicting stimulated peptide pools and resulting selected TCRs and accompanying CDR3 sequences. **B)** Frequency and culture origin of top 10 chosen TCR CDR3 sequences. Culture conditions include CD8^+^ T cells + autologous DCs, PBMCs + MHC-II blockade, and the original 11-day total PBMC culture.

**Supplemental Figure 9:**
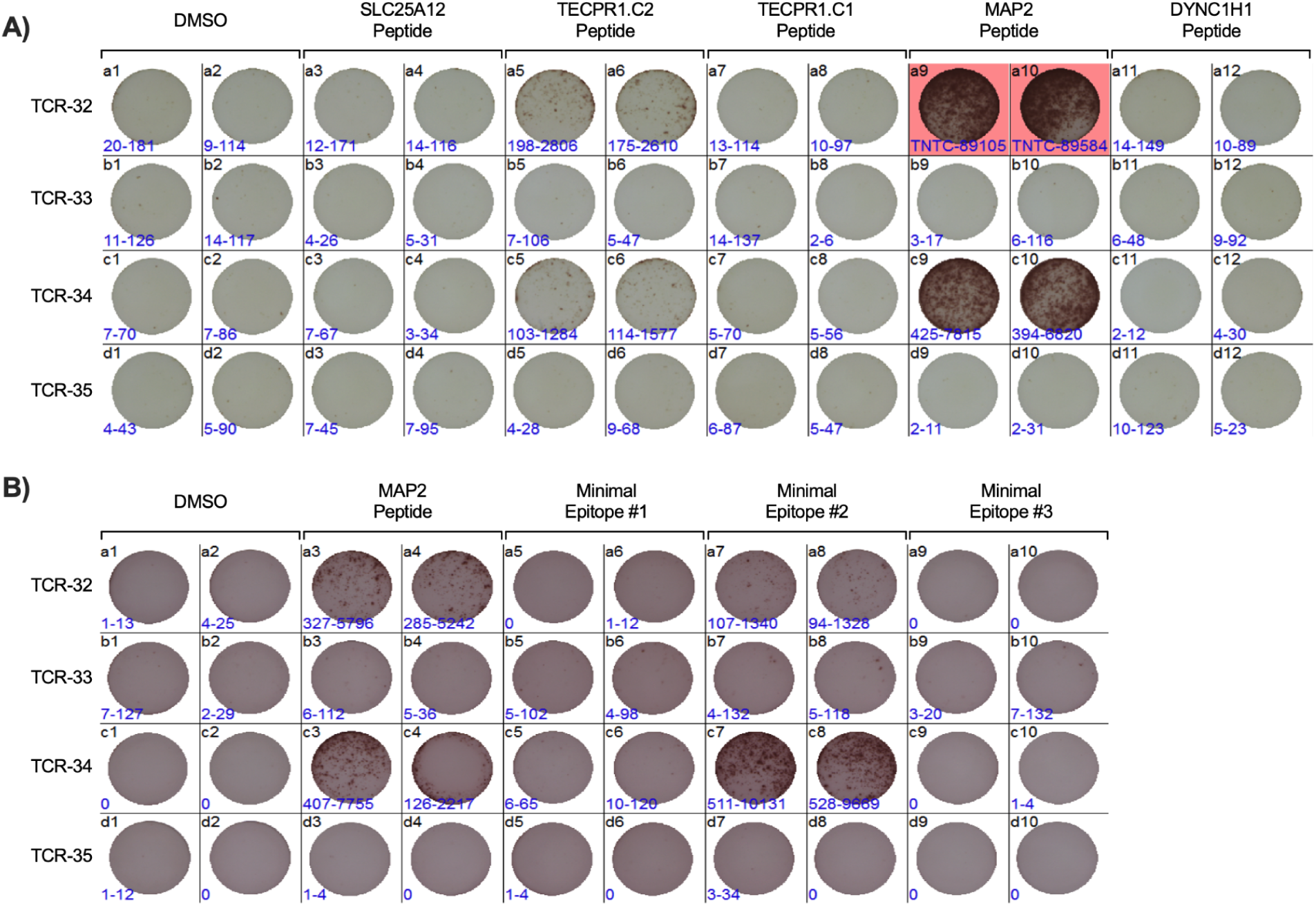
Supporting raw ELISPOT data assessing CD8-derived TCR-T specificity and minimal epitopes. **A)** Representative IFNγ ELISPOT images of chosen TCRs from Pool 2 against pooled peptides of origin; SFU per 50,000 T cells (n=2 technical replicates). Pools 1 and 3 (not pictured) contained only negatives. **B)** For all TCRs from pool 2, raw ELISPOT images comparing vaccine long peptides and corresponding predicted minimal epitopes; SFU per 50,000 T cells (n=2 technical replicates).

**Supplemental Table 1:**
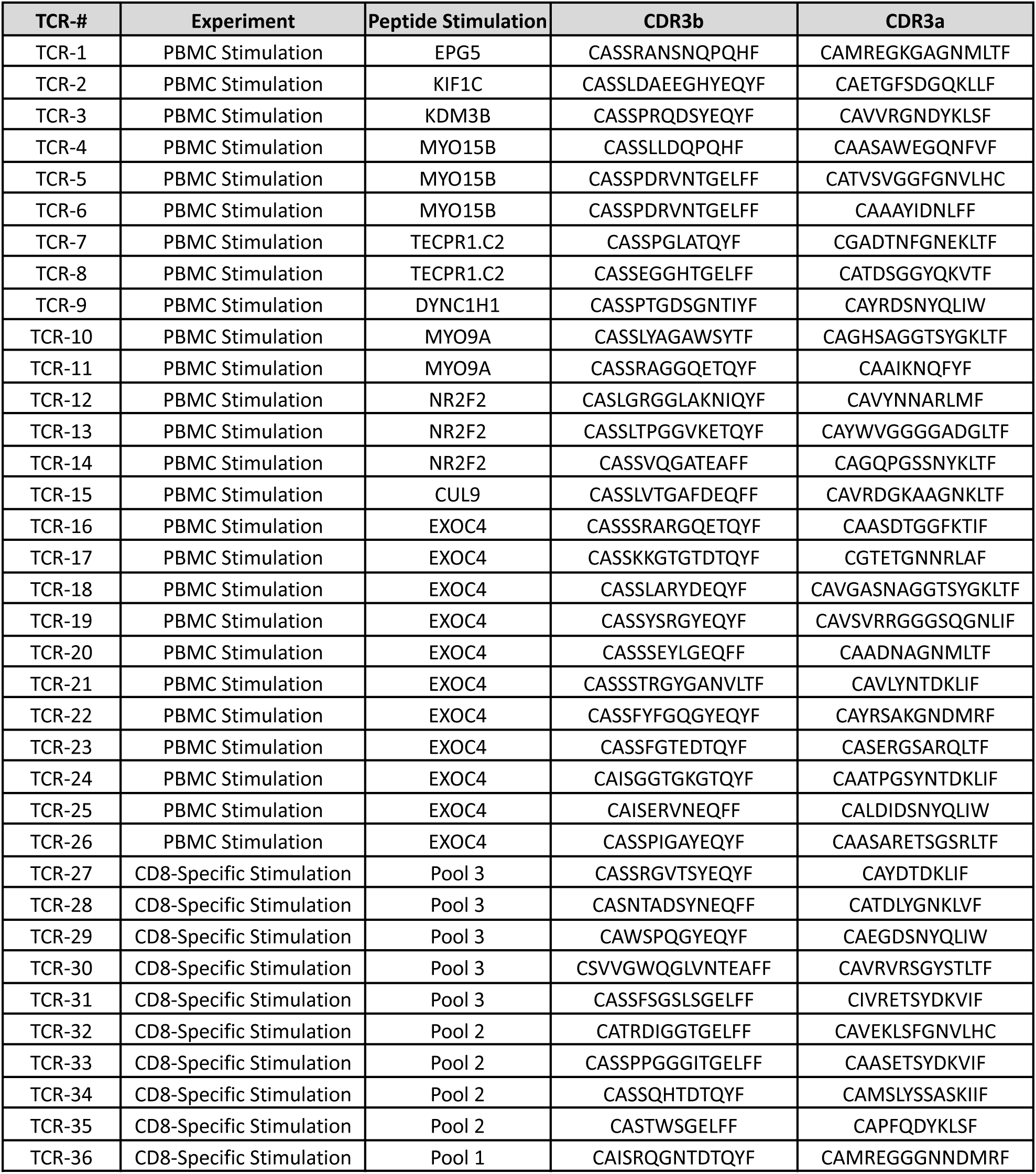
Tested TCR CDR3 sequences. CDR3 alpha and beta sequences for TCRs 1-26 (whole PBMC stimulation) and TCRs 27-26 (CD8-specific stimulation).

